# Identifying and testing marker-trait associations for growth and phenology in three pine species: implications for genomic prediction

**DOI:** 10.1101/2020.12.22.423987

**Authors:** Annika Perry, Witold Wachowiak, Joan Beaton, Glenn Iason, Joan Cottrell, Stephen Cavers

**Affiliations:** UK Centre for Ecology & Hydrology Edinburgh, Penicuik, Midlothian, EH26 0QB, UK; Institute of Environmental Biology, Faculty of Biology, Adam Mickiewicz University in Poznań, Poland; James Hutton Institute, Craigiebuckler, Aberdeen, AB15 8QH, UK; Northern Research Station, Forest Research, Roslin, EH25 9SY, UK

**Keywords:** Marker-trait association, predictive model, genetic variation, local adaptation, common garden trial, quantitative traits, SNP array, Scots pine, *Pinus mugo* complex

## Abstract

In tree species, genomic prediction offers the potential to forecast mature trait values in early growth stages, if robust marker-trait associations can be identified. Here we apply a novel multispecies approach using genotypes from a new genotyping array, based on 20,795 SNPs from three closely related pine species (*Pinus sylvestris, Pinus uncinata* and *Pinus mugo*), to test for associations with growth and phenology data from a common garden study. Predictive models constructed using significantly associated SNPs were then tested and applied to an independent multisite field trial of *P. sylvestris* and the capability to predict trait values was evaluated. One hundred and eighteen SNPs showed significant associations with the traits in the pine species. Common SNPs (MAF > 0.05) associated with bud set were only found in genes putatively involved in growth and development, whereas those associated with growth and budburst were also located in genes putatively involved in response to environment and, to a lesser extent, reproduction. At one of the two independent sites, the model we developed produced highly significant correlations between predicted values and observed height data (YA, height 2020: r = 0.376, *p* < 0.001). Predicted values estimated with our budburst model were weakly but positively correlated with duration of budburst at one of the sites (GS, 2015: r = 0.204, *p* = 0.034; 2018: r = 0.205, *p* = 0.034-0.037) and negatively associated with budburst timing at the other (YA: r = -0.202, *p* = 0.046). Genomic prediction resulted in the selection of sets of trees whose mean height was taller than the average for each site. Our results provide tentative support for the capability of prediction models to forecast trait values in trees, while highlighting the need for caution in applying them to trees grown in different environments.

## 1. Introduction

A primary goal of association genetics in long-lived organisms such as trees is to develop capacity to predict, at early life stages, the trait values of mature trees. However, the main traits of interest – such as height, volume, disease tolerance – are typically controlled by many genes, show quantitative variation, and may vary in expression and heritability depending on the environment in which they are assessed (Goddard and Hayes, 2009, Schlichting, 1986). Therefore, a high number of genetic markers screened in a large number of samples, which have also been accurately phenotyped, ideally in multiple environments, are required to develop robust predictive models for these traits. However, the power of genetic association studies is growing rapidly with improvements in the scale, accuracy and cost of high-throughput sequencing and genotyping. In particular, the accessibility of cost-effective high-throughput genotyping has benefited the study of non-model organisms, especially those for which genome assembly is challenging due to genome size and/or complexity (Prunier et al., 2016, Zimin et al., 2017). Allied to parallel efforts in building phenotype datasets, these technical and analytical advances mean association genetics in a range of tree species is now tractable.

In tree breeding, genome-wide single nucleotide polymorphism (SNP) markers can be used to predict breeding values and significantly increase the rate of gain in subsequent generations in a process known as genomic selection or genomic prediction (Meuwissen et al., 2001). The use of association analyses to identify SNPs significantly associated with traits of interest can further reduce and refine the number of SNPs used in predictive models. In this context, genomic prediction aims to increase the efficiency of breeding programmes to improve timber yield and quality and reduce losses due to pests and diseases in commercial forestry. Increasingly, it is also being used to screen natural populations for their adaptive potential to future threats such as climate change and disease (Isabel et al., 2020, Capblancq et al., 2020). To develop predictive models, multiple, ideally independent, trials are necessary to identify, test and validate the SNPs associated with each trait. In trying to apply genomic prediction approaches to populations outside breeding programmes, there are the additional challenges of comparative genetic complexity (Herbert et al., 1999), a lack of pedigree information, and an entirely different selection regime.

In the association analysis and prediction model development phase, groups of closely related species that are differently adapted but have similar genetic backgrounds can be useful experimental systems in which to search for parallel signatures of selection at the genomic level (Wachowiak et al., 2015). For such groups, a multispecies genomic approach can improve the power to detect genes involved in adaptation and show whether orthologous loci contribute to adaptive variation in different species (Neale and Ingvarsson, 2008). Multispecies approaches are reported to improve our understanding of both transcriptomes and genomes (for example, Ahrazem et al., 2019, Cornell et al., 2007, Leebens-Mack et al., 2019, Pellegrini et al., 1999, Polturak et al., 2018): the use of comparative species analyses provides a wide phenotypic base and shared evolutionary history for the identification of inter- and intra-specific genetic variation (van Kleunen et al., 2014). Making use of this multispecies approach to select SNPs for genomic prediction may also help to locate them in influential loci and potentially improve the generality of models based upon them.

Globally, pines are among the most important commercial tree species (Kanninen, 2010) and are ecosystem-defining in vast areas of forest across the northern hemisphere. Understanding the genetic architecture of key adaptive traits in pines, such as growth, form, disease resistance and phenology is of interest to a wide range of stakeholders, including the forestry industry and conservationists. Due to their large size and complexity, the assembly of pine genomes is particularly challenging, and has only been satisfactorily achieved for loblolly pine (*Pinus taeda*; Zimin et al., 2014) and sugar pine (*Pinus lambertiana*; Stevens et al., 2016), which are among the largest genomes ever sequenced and assembled. However, thousands of polymorphic regions potentially suitable for use in genotyping in pine species have already been discovered using high-throughput sequencing methods such as whole transcriptome studies (Blanca et al., 2012, Chancerel et al., 2011, Durán et al., 2019, Geraldes et al., 2011, Liu et al., 2014, Parchman et al., 2010, Trick et al., 2009, Wachowiak et al., 2015).

So far, most prediction models have been developed and tested for wood or fruit quality (Kumar et al., 2012, Minamikawa et al., 2017, Muranty et al., 2015, Beaulieu et al., 2014, Isik et al., 2016, Resende et al., 2012a, Resende et al., 2012b, Thistlethwaite et al., 2017), although a few have targeted disease resistance (Westbrook et al., 2020, Stocks et al., 2019). In pines, association studies and tests of genomic prediction have been performed for serotiny (*Pinus pinaster*, Budde et al., 2014; *Pinus contorta*, Parchman et al. 2012), circumference, height, stem straightness (*Pinus pinaster*, Bartholomé et al., 2016), oleoresin flow (*Pinus taeda*, Westbrook et al., 2013) and growth and wood quality traits (*Pinus sylvestris*, Calleja-Rodriguez et al., 2020).

Here we study three closely related pine species (*P. sylvestris, Pinus mugo* and *Pinus uncinata*) which have contrasting growth habits and are adapted to different environments. The species are members of the same monophyletic group within Pinaceae (Grotkopp et al., 2004), having diverged within the last 5 million years (Wachowiak et al., 2011) and have the same number of chromosomes (2n = 24). The three species have weak reproductive barriers between them and share many ancestral polymorphisms (Lewandowski et al., 2000; Wachowiak et al., 2013). We used trait data from a common garden glasshouse experiment (Wachowiak et al., 2018a) along with genotyping data from a large new multispecies *Pinus* SNP array (Perry et al., 2020) to identify SNPs associated with growth and phenology in the three species and to determine their putative function. A range of genomic prediction models were developed, and then tested using a subset of the *P. sylvestris*. Finally, we evaluated the potential of the models for genomic prediction, by testing the best performing set to estimate trait values in an independent multisite *P. sylvestris* field trial. We discuss the potential and limitations of the models for genomic prediction.

## 2. Methods

Experimental design and analyses performed in the study are summarised in Figure 1.

**Figure 1.**
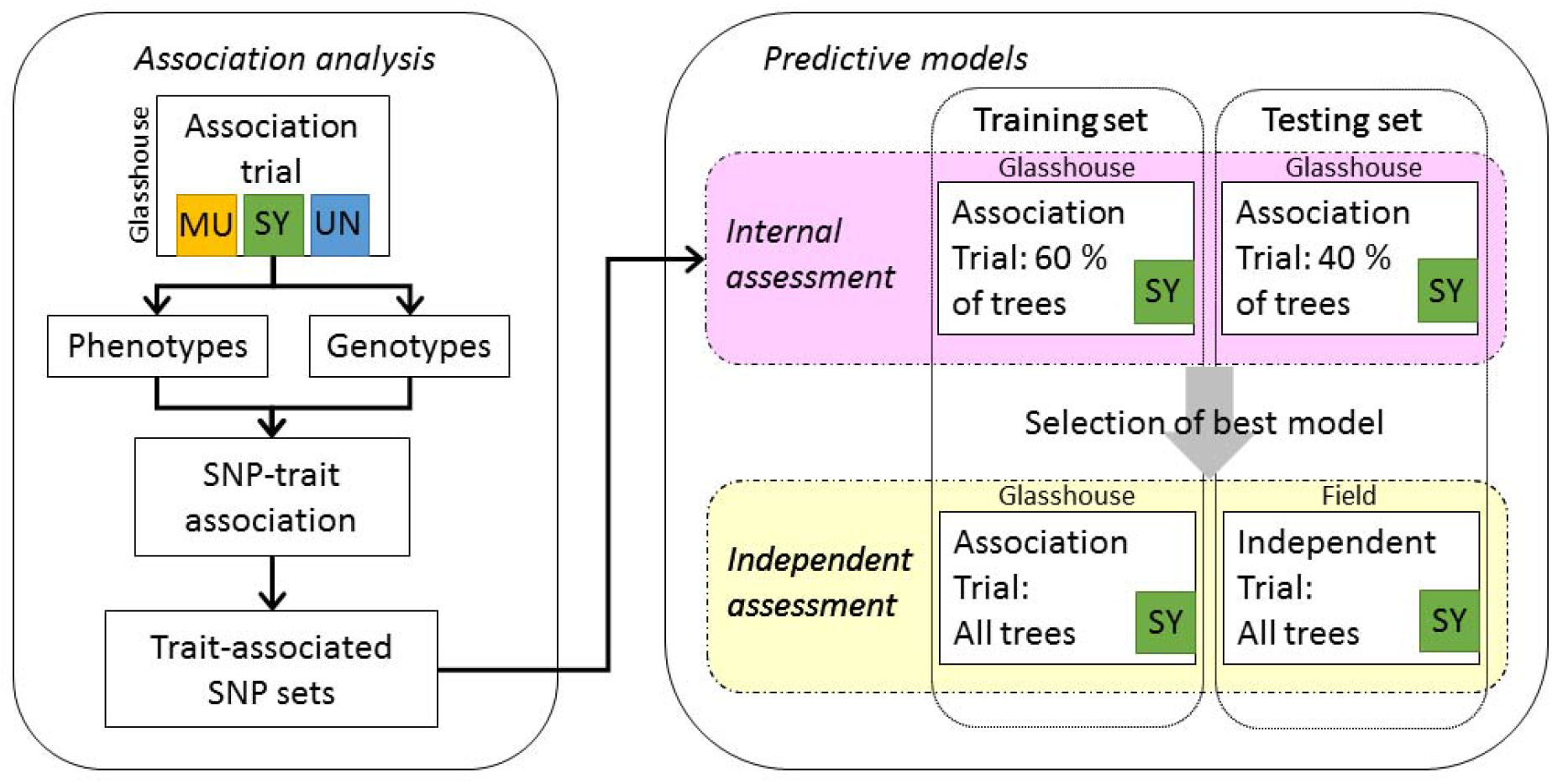
Plant material, datasets and analyses used in the study. MU: *P. mugo*; SY: *P. sylvestris*; UN: *P. uncinata*

### 2.1. Plant material and phenotype assessments

Collection of plant material, experimental design and phenotype assessments for the common garden glasshouse trial (referred to hereafter as the association trial) are described by Wachowiak et al., (2018a). Briefly, open-pollinated seeds of the three pine species were collected from three to five trees per population from twenty-eight natural populations in Europe covering the geographic range of each species (Figure 2). The collection consisted of thirteen populations of *P. sylvestris* (SY), nine *P. mugo* (MU), and six *P. uncinata* (UN). Seeds from each maternal tree were sown on trays of compost in spring 2010. After germination, a provenance–progeny trial was established in an unheated glasshouse at the UK Centre for Ecology and Hydrology, Edinburgh, UK (latitude 55.861261, longitude - 3.207819). Seedlings were grown under natural light with automatic watering applied during the growing season. The trial was divided into 25 randomized blocks with up to five families per population, of which the first 18 blocks were analysed by Wachowiak et al. (2018a). A summary of the counts of populations, families and total numbers of individuals are provided in Table S1. Phenology (traits assessed: BS, timing of bud set, BB, timing of budburst) and growth (traits assessed: H, total height; I, annual increment - the increase in height from one year to the next) were recorded for every seedling to evaluate within- and between-species variation (species means for trees sampled in this study recorded in Table S2).

**Figure 2.**
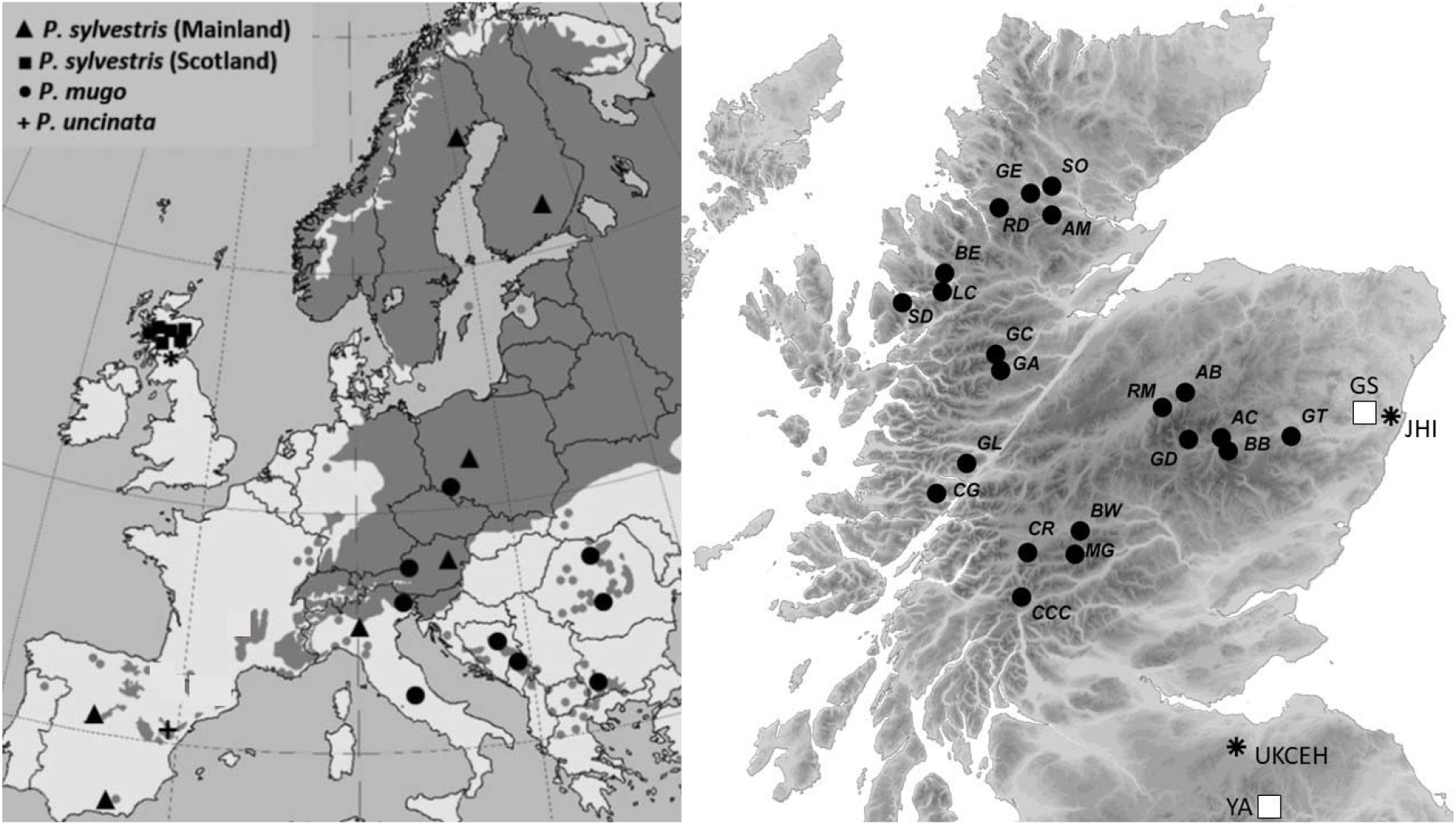
Geographic location of sampled pine populations across Europe (map on left: association trial) and Scotland (black circles in map on right: independent trial). Multi-site field trial locations (GS, Glensaugh; YA, Yair) are indicated with white squares on right hand map; glasshouses used to grow the association and independent trials (JHI, James Hutton Institute; UKCEH, UK Centre for Ecology and Hydrology) are indicated with asterisks on right hand map.

Bud set was defined as the time when a visible apical bud with clearly developed scales was formed at the tip of a stem in each seedling and was recorded as the number of days since the date on which the first plant to set a terminal bud in the trial was observed (in the first year of growth: BS2010). Budburst was scored when new needles emerged around the tip of the apical bud in the main stem and was measured as the number of days since the date on which the first plant to burst bud was observed (in the second and third years, BB2011, BB2012). Phenology observations were conducted twice a week. The height of all pines was measured annually from the second to fourth year of the pine growth (H2011, H2012, H2013). The annual increment was estimated for growth between 2011-12 (I2012) and 2012-13 (I2013). On the rare occasions that height was lower than in the previous year (due to, for example, human error or the loss of the leader) measurements were adjusted to ‘NA’. To assess the proportion of variation that is under genetic control, the narrow sense heritability (*h*^*2*^) and associated standard error for each trait was estimated using the GRM [genetic relationship matrix]-based restricted maximum likelihood (GREML) procedure implemented in GCTA (Yang et al 2011).

An independent multi-site, field-based provenance-progeny trial of *P. sylvestris* (referred to hereafter as the independent trial) was also phenotyped and genotyped using the same genotyping array and was used to test the predictive power of models developed using plants in the association trial described above. This trial was selected as there was commonality in the traits measured and geographic range of Scots pine populations used with the glasshouse trial, however the number of genotyped samples were considered to be too low to perform association analyses and it was therefore used instead to test the predictive power of the models in two distinct environments instead. Seeds from eight families from each of 21 native Scottish *P. sylvestris* populations (Figure 2) were collected in March 2007 and germinated at the James Hutton Institute, Aberdeen (latitude 57.133214, longitude -2.158764) in June 2007. A subset of trees from two of these sites were genotyped as part of this study: a site in the Borders of Scotland (Yair, YA: latitude 55.603625, longitude -2.893025) was planted in October 2012; a site in Aberdeenshire (Glensaugh, GS: latitude 56.893567, longitude - 2.535736) was planted in spring 2012. Trees transplanted to YA were initially grown in an unheated glasshouse whereas trees transplanted to GS were started in pots outside. The two transplantation sites also generally experience different climates, with the YA site typically warmer and drier than the GS site (Table S3) and with a longer growing season.

At each trial site, trees were planted in four randomised blocks at 3 m x 3 m spacing. A guard row of Scots pine was planted around the periphery of the blocks. Each block comprised one individual from each of eight families per 21 populations (168 trees). A summary of the counts of populations, families and total numbers of individuals are provided in Table S1. Budburst and height were assessed annually from 2015. Height was measured in the winter before the growing season began from 2015 to 2020. Height was also measured before the start of the second growing season in March 2008. The annual increment was estimated as the increase in height from one year to the next. Each tree was assessed for budburst stage annually from 2015 until 2019 at weekly intervals from early spring until budburst was complete. Seven distinct stages of budburst were defined (Table S4). The number of days for each tree to reach each stage of budburst, starting from the day the first tree was observed at each stage at each site, was recorded. When trees progressed through budburst stages rapidly, skipping a stage between assessments, a mean value was taken between the two assessment dates. The duration of the core stages of budburst (time taken to progress from stage 4 to stage 6) was also estimated. Although the method used to record budburst was not identical among the association trial and independent trial, the trait as described by Wachowiak et al., (2018a) is equivalent to stages 5 and 6 in the independent trial.

To better understand the relationship between different traits and for individual traits across different years, Pearson’s correlation coefficient and significance values were estimated for trees in the association trial for each species separately and for trees in the multi-site independent trial for each site separately using a package ‘Hmisc’ (Harrell Jr, 2020) in R (R Core Team, 2020). Data on individual traits measured over multiple years enabled their consistency among years to be assessed, as inter-annual variation may occur due to seasonal environmental variation, developmental variation and/or maternal effects (Vivas et al., 2020). Pearson’s correlation coefficient and associated significance values between budburst timing and duration among years and stages in the independent trial were also examined.

Nested ANOVA was performed for growth and phenology in the independent trial to assess within-site spatial heterogeneity for each site. Data for all trees in the trial was used (i.e. not just the subset of genotyped trees), with population as a fixed effect, and families nested within population and block as random effects.

### 2.2. Genotyping array

The design of the array, genotyping and SNP calling are as described by Perry et al., (2020). Briefly, an array comprising 49,829 single nucleotide polymorphisms (SNPs) was used to genotype 1,920 DNA samples (from needles of four pine species: the species included here plus *Pinus uliginosa*) according to the Affymetrix Axiom Assay protocol on a GeneTitan and following genotyping, genotype calls were performed using Axiom Analysis Suite as recommended by the manufacturer. A subset of trees from the association trial described in the previous section were genotyped and consisted of twelve populations of SY (N = 461) and five populations each of MU (N = 145) and UN (N = 201). Up to 10 trees were genotyped per family (except for population SY33 which was genotyped up to a maximum of 14 trees per family). Five families were genotyped per population with the exception of the following; SY44 (N families = 4), SY30 (N families = 3) and MU5 (N families = 3). Samples were filtered to remove all those with a call rate < 80 % (N removed: MU = 30; SY = 5; UN = 10). A summary of the counts of genotyped populations, families and total numbers of individuals are provided in Table S1.

The independent trial of *P. sylvestris* was also partially genotyped at each site: 100 trees from YA (15 % of the trees at the site) and 108 trees from GS (16 % of the trees at the site), each comprising the same five populations (Beinn Eighe, BE; Glen Affric, GA; Glen Loy, GL; Glen Tanar, GT; Rhidorroch, RD) with 19-22 individuals per population for each site. There were 7-8 families genotyped for each population with 1-3 half-siblings in each family at each site. These datasets are henceforth referred to as YA-SY and GS-SY. A summary of the counts of genotyped populations, families and total numbers of individuals are provided in Table S1.

### 2.3. Population genetic structure, kinship and statistical power

On the basis of the SNP genotyping results in the association trial, population genetic structure was assessed visually by constructing a neighbour joining tree in the R package ‘ape’ (Paradis and Schliep, 2019) based on a distance matrix generated in TASSEL version 5.2.39 (Bradbury et al., 2007). SNPs with call rate < 80 % (N = 48) were excluded. Pairwise kinship (centred identity by state) was estimated for each species independently using all polymorphic markers in TASSEL. The degree of skewness in the distribution within each species’ matrix was calculated using the D’Agostino skewness test in the R package ‘fBasics’ (Wuertz et al., 2020). The statistical power of each species’ dataset (MU; SY; UN), the *P. mugo* complex (MU-UN), and the full dataset including all species (MU-SY-UN) to detect true associations between SNPs and adaptive traits was estimated using the method reported by Wang and Xu (2019) under the following assumptions: nominal type 1 error (false positive) = 0.05; QTL size = 0.05. Statistical power was estimated at different levels of polygenic effect (λ): from 0.1 (where polygenic variance is 10 % of phenotypic variance) to 10 (where polygenic variance is 10 x phenotypic variance). Genotype frequencies of all SNPs subsequently found to be significantly associated with the adaptive traits in the MU-UN dataset were checked in each species separately (MU and UN) to assess the contribution of each species to associated genetic variation.

### 2.4. Genetic associations and putative functions

Using results from the association trial, identification of SNPs potentially associated with phenology (traits: budburst and bud set) and growth (traits: height and increment) was conducted for each trait in each year. For all analyses, raw phenotypic data were used. Association with SNPs was tested in each species separately (MU; SY; UN) as well as in all species combined (MU-SY-UN) and in the *P. mugo* complex (MU-UN). A mixed linear model (MLM) with a covariance (kinship: centred identity by state) matrices and matrices derived from principal component (PC) scores (separate matrices were constructed for each species/species set analysed), to allow for population stratification, among individuals was fitted to each locus independently in TASSEL (version 5.2.39). The proportion of true null hypotheses was estimated using a false discovery rate (FDR) approach, retaining SNPs associated with traits with adjusted *p* values < 0.05. Two association analyses were carried out, firstly including all polymorphic SNPs irrespective of minor allele frequency (MAF) value and secondly, by applying a MAF filter (excluding SNPs with MAF < 0.05).

A multi-locus mixed model (MLMM) approach, with 10 steps, was used to identify loci which have large effects (Segura et al., 2012). Highly significant SNPs (based on estimations of genetic variance, *p* < 0.001) were included in a forward-backward stepwise approach, one by one, as cofactors in the model. The kinship matrix used in the MLM approach was also included, but PC scores were not used. Rather than using PC scores to estimate population structure the MLMM approach uses a kinship matrix to describe the covariance structure, which is thought to perform better when population structure is complex (Segura et al., 2012). The multiple Bonferroni criterion, defined as the largest model whose cofactors all have a *p*-factor below a Bonferroni-corrected threshold of 0.05 (Dunn, 1961), was used to indicate the best model.

SNPs were divided into two classes on the basis of their minor allele frequency: MAF > 0.05: common; MAF < 0.05: rare. As the majority of traits are controlled by many genes of very small effect it is likely to be important to consider every SNP identified. Each SNP found to be significantly associated with a trait (when no MAF filter was applied, in order to compare the putative function of genes containing rare and common variants) was also examined to compare the putative function of the genes on which they are located with the trait in question. To do this, the full unigene sequence in which each SNP is located was BLASTed against the uniprotkb_viridiplantae database, the result with the highest score (minimum e-value 1E-50) for each unigene was retained, and the putative function determined by a literature survey using the search term ‘*protein name* function plant’. Where the protein was uncharacterised, the protein domain and/or family was recorded and the most likely function inferred. Where putative functions could be determined the genes were grouped according to their role in the following phenotypic responses: ‘Response to environment’ (including abiotic and biotic stress response), ‘Growth and development’ (including cell division, differentiation and senescence); ‘Reproduction’ (including flowering time and seed yield). Although many cellular processes (e.g. metabolism, signalling pathways, DNA binding, transcription, translation) were also identified as putative functions, these were assumed to be underlying control and expression of phenotypic functions and were not assigned a function.

### 2.5 Prediction models: construction and internal assessment

Phenotypic prediction multiple linear regression models were constructed in R using data generated from the association trial. A number of different models were constructed and compared using different sets of SNPs and different traits to train the model (the different models assessed are listed in Table 3). Predictive models were constructed using SNPs identified as potentially associated with variation in phenology (trait: budburst: BB2011) and growth (traits: height and increment: H2013 and I2013). To assess the relative contribution of SNPs identified using the multispecies compared to a single species approach, predictive models for both growth and phenology were constructed using SNPs identified from either a) all species’ datasets (i.e. MU-SY-UN, MU-UN and SY); b) just datasets containing SY (i.e. MU-SY-UN and SY); c) just SY. Predictive models were also constructed using the same number of randomly selected SNPs from all polymorphic loci with the same proportion of rare and common SNPs as the other prediction models. Ten sets of randomly selected SNPs were tested for each trait and 95 % confidence intervals were reported. Additionally, predictive models were constructed using all available polymorphic SNPs for SY. All models for both phenology and growth (SNPs associated with trait from MU-UN-SY, MU-UN and SY; SNPs associated with trait from MU-SY-UN and SY; SNPs associated with trait from SY; random SNPs; all polymorphic SNPs) were run both with and without a MAF filter (retaining only SNPs which were common in the datasets from which the significant associations were originally identified). As recommended when the number of loci is greater than the number of samples, we elected to construct the prediction model based on all polymorphic SNPs using ridge regression with the R package ‘rrBLUP’ (Endelman, 2011), rather than the multiple linear regression method. For all models, where necessary, family means were used to replace missing genotype data. The predictive models were initially run using an internal training set comprising 60 % of SY trees from the association trial, which had been used to identify associated SNPs, and predictive accuracy assessed using the remaining 40 % of SY trees (the internal testing set) in the association trial. Models were run using SY trees and not UN or MU as our intention was to carry out subsequent testing of the models in this species alone. We used budburst and growth but not bud set data as subsequent model testing was applied to data from an independent trial for which only these traits were available. Pearson’s correlation coefficient and significance for correlations between predicted values generated by the predictive models and observed values for both phenology and growth (both H2013 and I2013 were tested to see which performed best for the growth model) were estimated using the R package ‘Hmisc’ (Harrell Jr, 2020).

SNPs used in each prediction model were assessed for their variation among SY populations using the R package ‘hierfstat’ (Goudet and Jombart, 2020). Basic statistics including overall observed heterozygosity (H_O_), mean gene diversities within populations (H_S_), inbreeding coefficient (F_IS_) and population differentiation (F_ST_) were estimated for each set of SNPs described above.

### 2.6 Prediction models: independent assessment

SNPs identified as potentially associated with budburst and growth were tested for their predictive power using genotype and phenotype data from an independent trial of *P. sylvestris*, established at contrasting sites (YA and GS) in 2012. Genotyped trees from YA and GS were assigned predicted values for both phenology and growth using multiple linear regression models constructed using either all available SNPs or only those found to be significantly associated with the trait (final predictive models, chosen based on their performance in the initial internal test). As models tend to work best in material that is closely related to those used in model development (Beaulieu et al 2011), the model using all available SNPs was also trained using only SY trees from Scotland grown in the association trial (N = 227). Observed values for growth (height and increment) and budburst (timing and duration) at multiple years (2015-2020 for increment, 2015-2019 for budburst) were compared with values generated by the predictive models. Multiple years were used to ensure that annual variation caused by seasonal differences could also be considered. Height is a cumulative measure, and therefore, only the most recent (2020) and the measurements made after the first year of growth (2008) were compared with the predicted values. Furthermore, the use of height measurements at both young and more mature ages allowed the impact of maternal effects to be examined and tested. In order to identify SNPs which are good predictors of final height, the use of trees whose traits are not confounded by maternal effects is important. To assess the performance of the predictive models, the Pearson’s correlation coefficient and significance values between predicted and observed values for phenology and growth were estimated for each site (GS and YA) separately using the R package ‘Hmisc’ (Harrell Jr, 2020). The use of two sites in independent testing also allowed comparison of the performance of the predictive models in different environments.

The effectiveness of using the predictive model as a genomic selection tool was also tested and compared with other selection methods. For each method, 10 trees were selected from each trial site: for genomic selection, the 10 trees at each site with the highest values generated by the predictive model were chosen; for phenotype selection, the 10 tallest trees at each site prior to the start of the second growing season (measured in March 2008) were chosen. The average height at 13 years (2020) of the 10 trees selected using each method was compared. The trees selected using each method were also compared to the 10 tallest trees at each site at age 13.

## 3. Results

### 3.1. Intra- and inter-specific trait variation

Bud set was, on average, earliest for MU and latest for SY with a mean difference of nearly 19 days between the two species (Table S2, Table S5). Bud set for UN occurred, on average, 8.28 days after MU and 10.63 days before SY. Budburst was similarly earliest for MU but was latest for UN in both years assessed although the mean difference between species was greater in 2012 (15.17 days) than in 2011 (5.42 days). For all years, on average, MU were the shortest trees and SY were the tallest with increment similarly greater in SY than in UN or MU. By 2013, SY trees were on average over double the height of the average MU tree, with UN trees on average just over two-thirds the height of the average SY tree. Narrow sense heritability estimates were highest for bud set (mean *h*^*2*^ = 0.78; Table S6) and height (mean *h*^*2*^ = 0.70) and lowest for increment (mean *h*^*2*^ = 0.44). Narrow sense heritability estimates were highest for SY (mean *h*^*2*^ = 0.71) and lowest for all species combined (mean *h*^*2*^ = 0.55).

Phenotypes for the independent trial are provided in Table S7. The relationships between duration and timing of each stage of budburst in the independent trial were examined. Due to missing data, only budburst stages 4 to 6 were analysed. Timing (time taken to reach each stage) showed a significant negative correlation with duration (time taken to progress from stage 4 to 6) of budburst at each year assessed for stage 4, but the relationship was positively correlated for stage 6 (Table S8). In contrast, the time to reach stage 6 showed a significant positive correlation with the duration of budburst. Time to reach stage 5 was both positively (GS) and negatively (YA) correlated with the duration of budburst. Budburst stages were highly positively correlated with one another in all years at both sites (Table S8). Therefore, stage 6 is used to represent timing of budburst for all further analyses.

The relationships among the traits measured over multiple years (including both correlations of individual traits over multiple years and correlations of different traits with one another) were examined in both the association trial (Table S9) and independent trial (Table S10). For MU and UN in the association trial, height and increment were highly significantly correlated over each year, as was budburst in each year. Although highly significant in all cases, the correlation coefficient associated with height in the first year of growth (H2011) reduced in each subsequent year for all three species (H2011 and H2012: MU, 0.74; SY, 0.81; UN, 0.71. H2011 and H2013: MU, 0.67; SY, 0.73; UN, 0.66). Bud set in 2010 was significantly negatively correlated with budburst in 2011 in both MU and UN but was significantly positively correlated with budburst in both 2011 and 2012 in SY. Furthermore, bud set in SY was also highly significantly correlated with height in all years and increment in 2012 and increment in 2013 was not significantly correlated with height in 2011 or 2012. In the independent trial, height in 2020 was highly significantly correlated with increment for all years in both GS and YA, whereas height in 2008 was not significantly correlated with height in 2020, as would be expected if maternal effects were expressed in the first year of growth, or increment in any year except 2017 in GS. There was little correlation among growth and phenology traits in either independent trial site except budburst timing in 2016 which was negatively correlated with height and increment after 2015 in YA and positively correlated with height in 2020 and increment in 2015 and 2016 in GS.

The block effect was low at the GS site of the independent trial (Table S11) and was only significant for budburst timing and duration in 2017 and for increment in 2020. In contrast, blocks were significantly different for all traits for the majority of years at YA. Budburst timing was not significantly different among populations at either GS or YA in any year and was only significant for budburst duration in 2018 (both sites) and 2016 (GS). There were significant differences among populations for height and increment for all years in GS but not for increment in 2016 or 2017 at YA. There were significant differences among families for all traits at both GS and YA but this was not the case in all years.

### 3.2. Summary of genotyping array

High quality genotypes (call rate > 80 %) were obtained for over 94 % of trees genotyped within the association trial (N = 762: MU, N = 115; SY, N = 456; UN, N = 191, Table S5). There were 9,583 high quality (call rate > 80 %) polymorphic SNPs which were shared among the three species (Table 1), with a further 1,352 SNPs which were polymorphic in at least two species and monomorphic in a third. SNP genotypes obtained for trees from YA and GS were all high quality (Table S7).

**Table 1.**
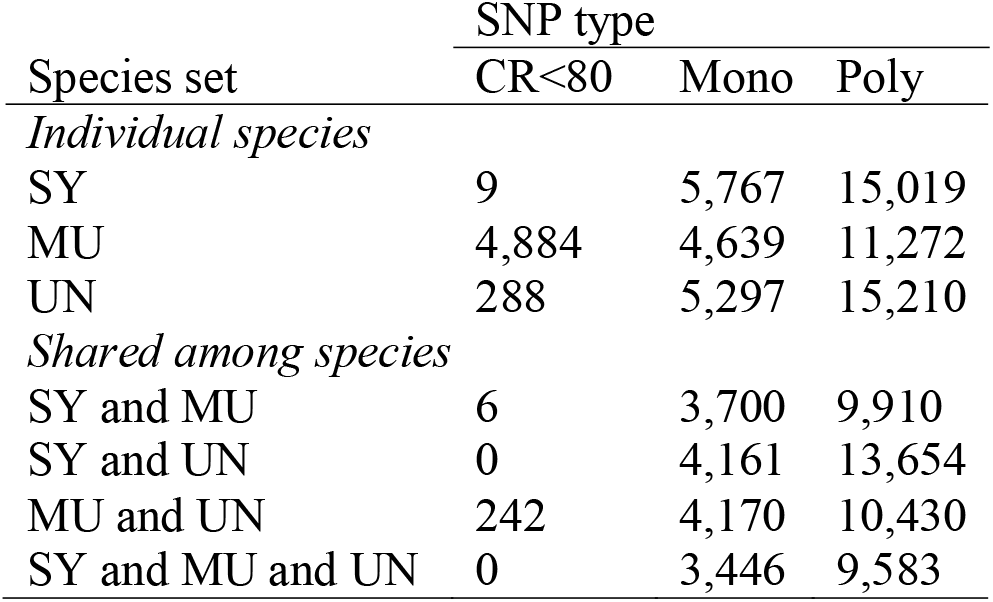
Counts for each type of SNP in individual species and shared among species. Species: SY, *P. sylvestris*; MU, *P. mugo*; UN, *P. uncinata*. SNP type: CR<80, call rate < 80 %; Mono, monomorphic; Poly, polymorphic.

### 3.3. Population genetic structure, kinship and statistical power

The neighbour joining tree generated from the distance matrix indicated the majority of the structure is among species and families with weak population structure, as reported in previous studies (Wachowiak et al., 2013, Wachowiak et al., 2018b). The pairwise kinship distribution was strongly skewed toward positive kinship values for each species (D’Agostino’s skewness test, MU: z = 101.389, *p* < 2.2 × 10^−16^; SY: z = 446.904, *p* < 2.2 × 10^− 16^; UN: z = 153.664, *p* < 2.2 × 10^−16^), as expected given the presence of half siblings in the association trial. These results support the use of mixed model approaches and the correction for population stratification prior to testing for genetic association.

The statistical power to detect true associations between SNPs and adaptive traits was found to be extremely low for both MU and UN even when the polygenic effect was assumed to be 10 x that of the phenotypic variance (Table S12). This is likely to be due to the low sample numbers, a conclusion supported by the result that the statistical power of SY was similarly low if the sample numbers were reduced to those of MU and UN: the statistical power remained low even when the polygenic effect was increased. The model based on the SY dataset was found to have relatively high statistical power, and the model based on the joint MU-UN dataset had lower power, but significantly more than for each species individually. The statistical power of a model based on the dataset that included all three pine species was found to be very high regardless of the polygenic effect. For these reasons, the following datasets were analysed for associations with traits: the *P. mugo* complex (MU-UN), SY, and all three pine species (MU-SY-UN).

### 3.4. Identification of loci associated with traits

One hundred and eighteen SNPs were identified as associated with phenology and growth in the three pine species (Table 2; Table S13) and included SNPs which were identified in more than one species’ datasets. There was very little overlap of individual SNPs associated among multiple traits or years: four SNPs were associated with more than one trait, of which only one (comp51128_c0_seq1_1529) was associated with both phenology (trait: BB2011) and growth (trait: I2013). The vast majority of SNPs were identified using the MLM approach (N SNPs = 113) rather than the MLMM approach (N SNPs = 14). There were nine SNPs identified as significantly associated with traits in both MLM and MLMM. Almost twice as many SNPs were significantly associated with phenology traits (N = 77) than growth traits (N = 42). Significantly associated SNPs were identified for all traits in all years except BB2012. The traits with the highest number of associated SNPs were BB2011 (N = 58), I2013 (N = 34) and BS2010 (N = 19), whereas other years/traits all had low numbers of associated SNPs (H2011, N = 3; H2012, N = 1; I2012, N = 1).

**Table 2.** Total number of SNPs associated with phenology and growth traits in the three pine species identified from a mixed linear model (MLM) in TASSEL and a multi-locus mixed model (MLMM) in R. SNPs identified in analyses with a minor allele frequency (MAF) filter (excluding MAF < 0.05) are in parentheses: common SNPs identified both with and without a MAF filter are to the left of the forward slash; SNPs identified only with a MAF filter are to the right of the forward slash)

| Trait | Species | MLM |  | MLMM |  |
| --- | --- | --- | --- | --- | --- |
|  |  | Common | Rare | Common | Rare |
| *Phenology* |  |  |  |  |  |
| BB2011 | MU-UN | 9 (3 /1) | 25 |  | 4 |
|  | SY | 7 (6/2) | 11 |  | 3 |
|  | MU-SY-UN | 4 (2/1) | 19 |  | 3 |
| BS2010 | SY | 4 | 14 |  |  |
|  | MU-SY-UN | (0/1) |  |  |  |
| *Growth* |  |  |  |  |  |
| H2011 | SY |  | 1 |  |  |
|  | MU-SY-UN | 1 | 1 |  |  |
| H2012 | MU-SY-UN | 1 (1/0) |  |  |  |
| H2013 | SY | 2 |  |  |  |
|  | MU-SY-UN | 4 |  | 1 |  |
| I2012 | MU-SY-UN | 1 |  |  |  |
| I2013 | MU-UN | 6 (5/0) | 1 |  | 1 |
|  | SY | 2 (1/0) | 20 |  | 4 |
|  | MU-SY-UN | 6 (3/0) | 4 | 1 | 1 |

Species codes: MU, *P. mugo*; SY, *P. sylvestris*; UN, *P. uncinata*; Trait codes: budburst (BB); bud set (BS); height (H); increment (I). Common: SNPs with MAF > 0.05; Rare: SNPs with MAF < 0.05

**Table 3.**
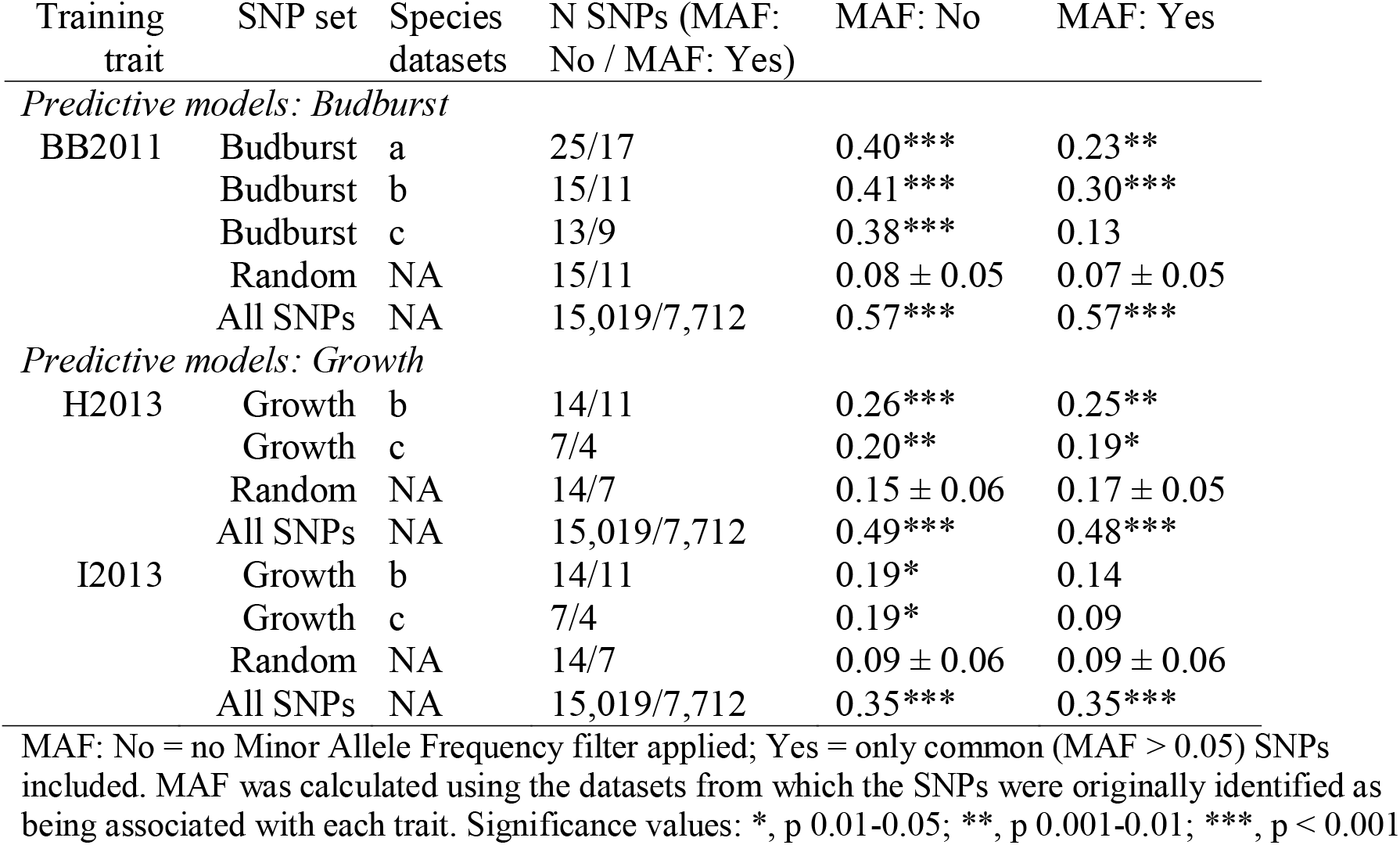
Pearson’s correlation coefficient I and associated significance values for comparison of predicted and actual values for each trait both with and without a MAF filter when using prediction models constructed with SNPs significantly associated with each trait (Budburst; Growth), random sets of SNPs (10 sets of randomly selected SNPs for each model with 95 % confidence intervals reported) or all polymorphic SNPs. Species datasets, SNPs identified as significantly associated with the trait in: a) all species’ datasets (i.e. MU-SY-UN, MU-UN and SY); b) just datasets containing SY (i.e. MU-SY-UN and SY); c) just SY. All models trained using a subset of the SY dataset and validated using the remaining SY trees.

| Training trait | SNP set | Species datasets | N SNPs (MAF: No / MAF: Yes) | MAF: No | MAF: Yes |
| --- | --- | --- | --- | --- | --- |
| *Predictive models: Budburst* |  |  |  |  |  |
| BB2011 | Budburst | a | 25/17 | 0.40*** | 0.23** |
|  | Budburst | b | 15/11 | 0.41*** | 0.30*** |
|  | Budburst | c | 13/9 | 0.38*** | 0.13 |
|  | Random | NA | 15/11 | 0.08 ± 0.05 | 0.07 ± 0.05 |
|  | All SNPs | NA | 15,019/7,712 | 0.57*** | 0.57*** |
| *Predictive models: Growth* |  |  |  |  |  |
| H2013 | Growth | b | 14/11 | 0.26*** | 0.25** |
|  | Growth | c | 7/4 | 0.20** | 0.19* |
|  | Random | NA | 14/7 | 0.15 ± 0.06 | 0.17 ± 0.05 |
|  | All SNPs | NA | 15,019/7,712 | 0.49*** | 0.48*** |
| I2013 | Growth | b | 14/11 | 0.19* | 0.14 |
|  | Growth | c | 7/4 | 0.19* | 0.09 |
|  | Random | NA | 14/7 | 0.09 ± 0.06 | 0.09 ± 0.06 |
|  | All SNPs | NA | 15,019/7,712 | 0.35*** | 0.35*** |

MAF: No = no Minor Allele Frequency filter applied; Yes = only common (MAF > 0.05) SNPs included. MAF was calculated using the datasets from which the SNPs were originally identified as being associated with each trait. Significance values: \*, p 0.01-0.05; \*\*, p 0.001-0.01; \*\*\*, p < 0.001

A higher number of SNPs associated with traits were identified in SY (N = 64) than in MU-UN (N = 44). Only one SNP (comp51128_c0_seq1_1529) was identified as significant in both datasets although it was associated with phenology (BB2011 for MU-UN) and growth (I2013 for SY): it was common in MU-UN but rare in SY (Table S13). A further 44 SNPs were found to be associated with traits when all species were combined within a single analysis, although 11 of these were also identified in SY and 23 were identified in MU-UN. Applying a multispecies approach led to the identification of 54 SNPs which would not have been identified if only the SY dataset had been used. When no MAF filter was applied prior to screening SNPs for association with the traits of interest, 37 SNPs were found to be common in at least one dataset. Applying a MAF filter identified a further five SNPs, all in phenology traits (Table 2), but also failed to identify 26 of the common SNPs identified when no MAF filter was applied.

Genotype frequencies for SNPs identified as significantly associated with adaptive traits in MU-UN were compared for UN and MU separately (Table S14). Diversity was much lower in UN than MU for the majority of SNPs: 23 of the 36 SNPs identified as associated with BB2011 were monomorphic in UN. In contrast, diversity in UN was much higher for SNPs identified as significantly associated with I2013 (Table S14). Similarly, the standard error for MU was more than twice that of UN for BB2011 (MU: 0.73; UN: 0.33) whereas the standard error for both species was similar for I2013 (MU: 0.47; UN: 0.32; Table S2).

### 3.5. Putative function of genes containing SNPs associated with traits

One hundred and eighteen SNPs associated with phenology and growth in the three pine species were located at 114 gene loci (two unigenes, comp48223_c0_seq1 and comp47733_c0_seq1, contained three SNPs each). One locus was originally identified in *Pinus radiata* (Doth_comp54682_c0_seq1_159), the remaining were identified following transcriptome sequencing in *P. sylvestris* and the taxa of the *P. mugo* complex (Perry et al., 2020: Table S13). The genetic sequences containing loci associated with each trait were found to be highly similar to proteins with a range of putative functions (Tables S17a-c). Of the SNPs identified when no MAF filter was applied, the majority of SNPs associated with bud set (all identified in SY) were found in genes that code for proteins putatively involved in growth and development (61.11 %) with a few (exclusively rare) SNPs found in proteins putatively involved in response to environment (22.22 %, Figure 3). In contrast, budburst had high numbers of associated SNPs (both rare and common) in genes that code for proteins putatively involved in response to environment and growth and development (mean contribution of putative function groups coded by genes containing SNPs significantly associated with budburst across species’ datasets as a percentage of the total number of proteins: 39.01 % and 39.09 % for growth and development and response to environment, respectively). Whereas the majority of SNPs associated with height were found in proteins putatively associated with growth and development, SNPs associated with increment were found in proteins putatively associated with both growth and development and response to environment. There are some differences among species in the putative function of proteins containing significantly associated SNPs: the majority of SNPs in SY are found in genes coding for proteins putatively associated with growth and development for all traits (Figure 3) whereas MU-SY-UN and MU-UN have higher proportions of SNPs in genes coding for proteins putatively associated with response to environment as well as growth and development. Of the five SNPs that were only identified as associated with a gene when a MAF filter was applied, one was putatively associated with response to environment, one with growth and development, one with all three functions and two with none of these functions (Table S15a-c).

**Figure 3.**
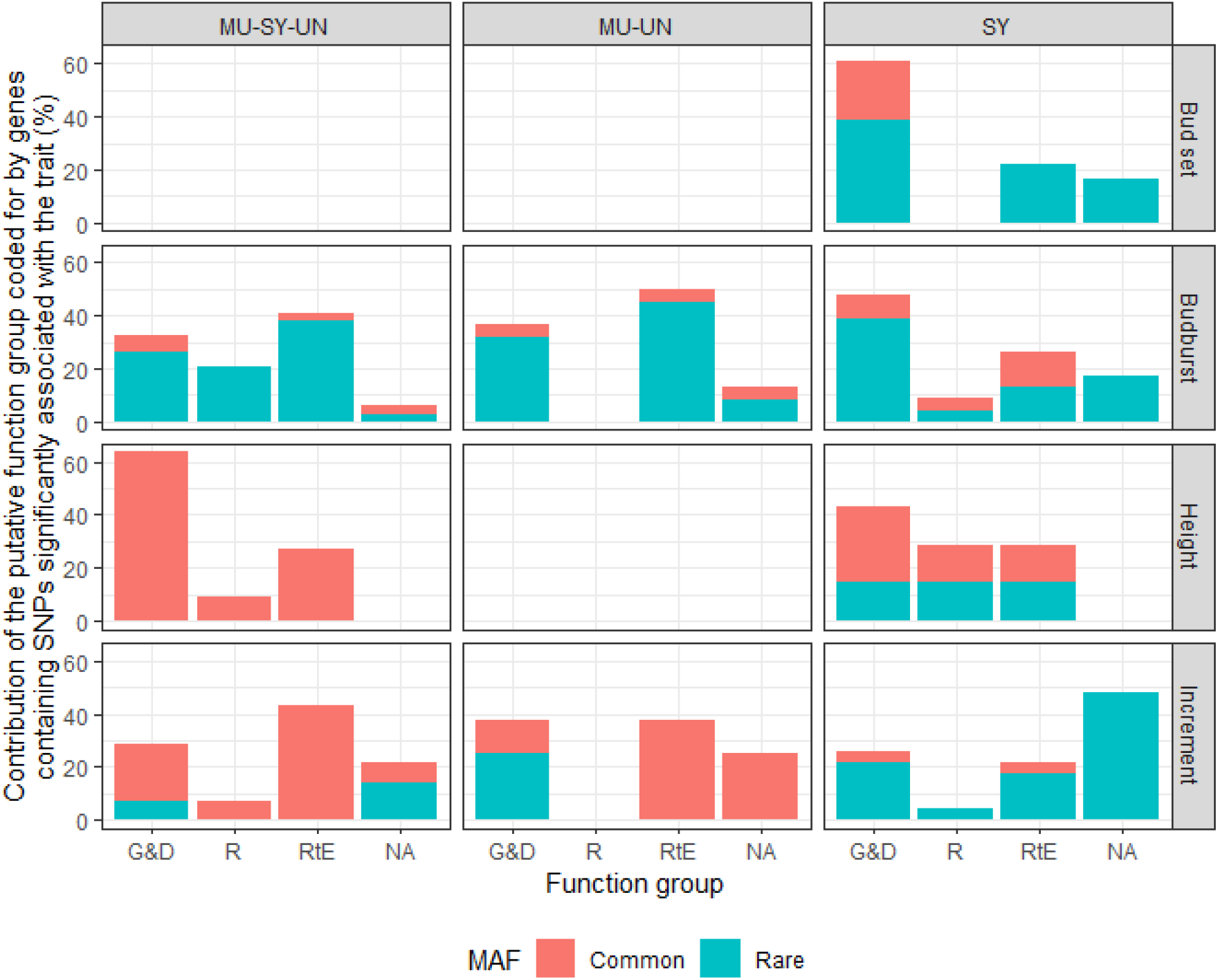
Contribution of putative function groups (G&D: growth and development; R: reproduction; RtE: response to environment) coded for by genes containing SNPs significantly associated with each trait (bud set, budburst, height and increment) identified when no MAF filter was applied and as a percentage of the total number of proteins identified for each trait for each species’ dataset (MU: *P. mugo*; SY: *P. sylvestris*; UN: *P. uncinata*). Proteins which were uncharacterised, for which no known function in plants was found or for which only cellular processes could be identified are categorised “NA”. Total for each trait may be higher than 100 % as there may be more than one putative function assigned to a single protein. MAF: minor allele frequency (MAF > 0.05: common; MAF < 0.05: rare)

### 3.6. Prediction models: construction and internal assessment

There was a large dropout in the number of SNPs which were suitable for inclusion in subsequent predictive models: of the 38 SNPs identified as potentially associated with growth (H2013 and I2013) in SY, MU-UN and MU-SY-UN datasets, 24 were monomorphic in either (or both) the SY and the independent *P. sylvestris* datasets (YA-SY and GS-SY). Therefore, 14 SNPs (nine associated with I2013, four with H2013 and one with both I2013 and H2013) were included in the model, of which eight were rare in the SY dataset (although only three were rare in the MU-SY-UN datasets in which they had been identified as associated with the traits). Of these 14 SNPs, five were identified in the SY dataset, four in the MU-SY-UN dataset, three in both the MU-UN and MU-SY-UN datasets and two in both the SY and MU-SY-UN datasets. For the predictive models for budburst, all SNPs significantly associated with BB2011 in the SY, MU-UN and MU-SY-UN datasets (N = 58) were considered for inclusion. Thirty three SNPs significantly associated in at least one of the association trial datasets were monomorphic in at least one of the SY, YA-SY and GS-SY datasets. The remaining 25 SNPs (13 were identified in SY or in both SY and MU-SY-UN; 11 were identified in MU-UN or in both MU-UN and MU-SY-UN; one was only identified in MU-SY-UN) were used to construct the predictive models for budburst. Eight of the SNPs were rare in the dataset in which they were identified. The SNPs used to construct predictive models for growth were found to have lower differentiation among populations (F_ST_ = 0.03 to 0.04, Table S16) than the full set of polymorphic SNPs for SY (F_ST_ = 0.06). The inbreeding coefficient (F_IS_) was -0.6 to -0.7 for the majority of SNP sets (Table S16) with a slightly higher value observed in the SNP set for the growth model using trait associated SNPs identified in the SY dataset (F_IS_ = 0.9). Observed heterozygosity and gene diversity (H_O_ and H_S,_ respectively) were both higher in the sets of SNPs which were filtered to include only those which were common in the original dataset.

The performances of each predictive model when tested internally using the association trial material (i.e. the strength and significance of the correlation of predicted values with the observed values for each trait) are summarised in Table 3. Models constructed using random SNPs were not successful in predicting values that were correlated with observed values for each trait, although the mean strength of the correlation was much higher for the random models trained using H2013 than for either I2013 or BB2011. For all models, except those constructed using random SNPs, those without a MAF filter always performed better than the equivalent models constructed using only common SNPs, although there was little difference in performance for models constructed using all polymorphic SNPs (Table 3). The predictive model for budburst constructed using SNPs which were identified in both SY and MU-SY-UN resulted in a slightly improved predictive ability (r = 0.41, *p* < 0.001; Table 3) compared to the models which included only SNPs identified in SY (r = 0.38, *p* < 0.001) or those identified in all species’ datasets (r = 0.40, *p* < 0.001). For these reasons, the final predictive model for budburst was constructed using SNPs identified in both MU-SY-UN and SY, with no MAF filter applied to the SNPs. The predictive model for growth also performed best when using SNPs identified in multispecies’ datasets (MU-SY-UN and SY: r = 0.26, *p* < 0.001; SY only: r = 0.20, *p* = 0.008). There were no SNPs associated with growth and identified exclusively in MU-UN which were also polymorphic in SY, YA-SY and GS-SY. Using H2013 as a training trait, the predictive model for growth performed more poorly using SNPs identified in the SY dataset than using SNPs identified in both the SY and MU-SY-UN datasets. However, with I2013 as a training trait in the same model, there was no difference in performance when the different SNP sets were used. There were highly significant positive correlations between observed H2013 and predicted values when using the predictive models for growth whereas using I2013 as the training trait for the predictive model resulted in far lower levels of correlation between predicted and observed values. Therefore, the final predictive model for growth, constructed using SNPs identified in both the SY and MU-SY-UN datasets with no MAF filter and using H2013 as a training trait, was chosen to be tested independently.

The effect of the trait used to train the model was also seen in comparisons of the performance of the models constructed using all polymorphic SNPs: for each trait, predicted values were more closely correlated with the observed values in models using budburst than in those using growth traits (H2013, I2013). Of the traits used to identify associated SNPs and construct the predictive models, the one with the lowest *h*^*2*^ (I2013) also had the lowest predictive ability in the SY dataset, whereas the trait with the highest *h*^*2*^ (BB2011) had the highest predictive ability.

### 3.7. Prediction models: independent assessment

Predicted values were estimated using the final predictive models for budburst and growth as well as models constructed using all available SNPs and compared with values observed in the independent trial. The independent field sites share populations and families but experience contrasting climates, allowing the models to be independently tested on traits measured in different environments. The predicted values for each trait were not significantly correlated with the observed values when using models constructed with all available SNPs when trained using the full SY dataset and only for increment in 2016 at GS when only SY from Scotland is used to train the model (Table 4). In contrast, a number of significant correlations were observed in the independent trial when using final predictive models for growth and budburst. The predicted values for budburst were found to be significantly positively correlated with the duration of budburst at GS in 2015 and 2018 (Table 4) indicating a possible effect of annual environmental variation on the predictive power of the model. They were also negatively associated with budburst timing at YA in 2017.

**Table 4.**
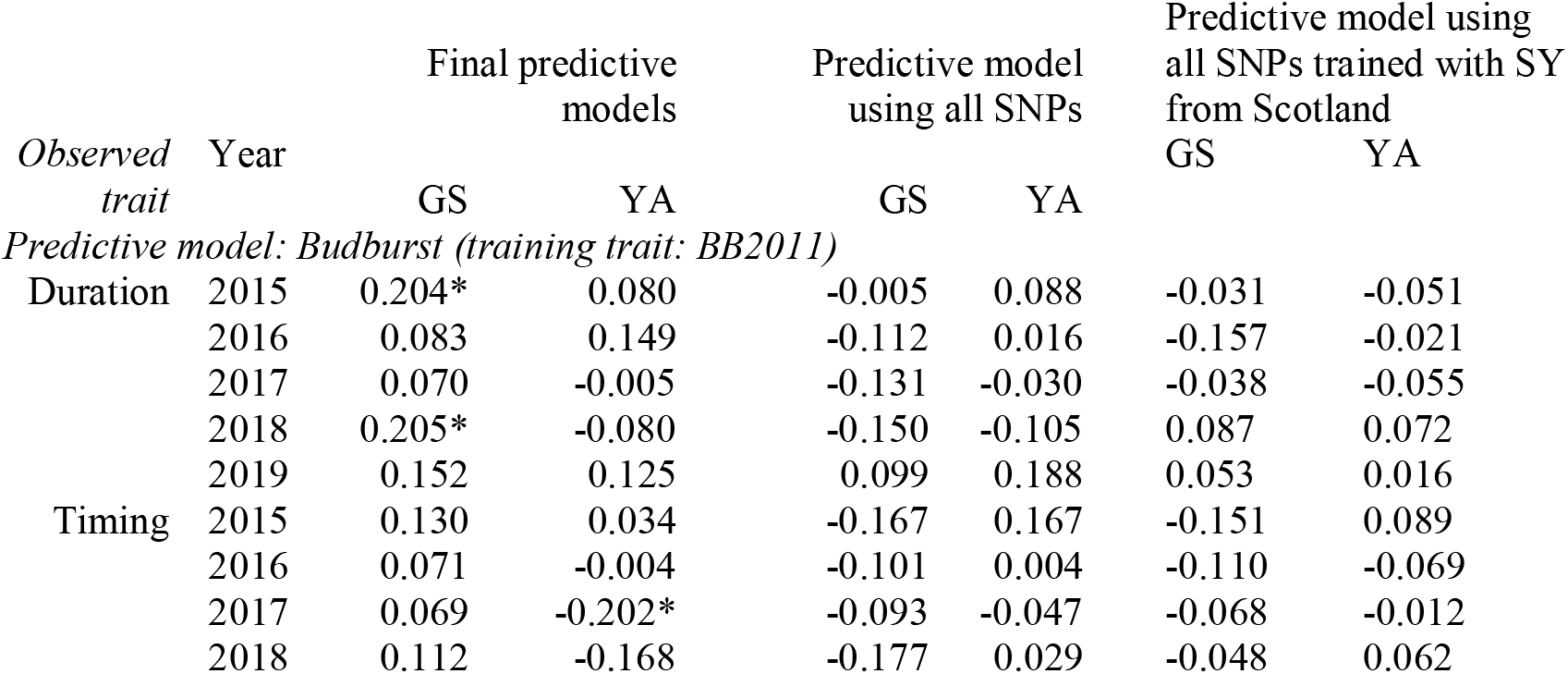
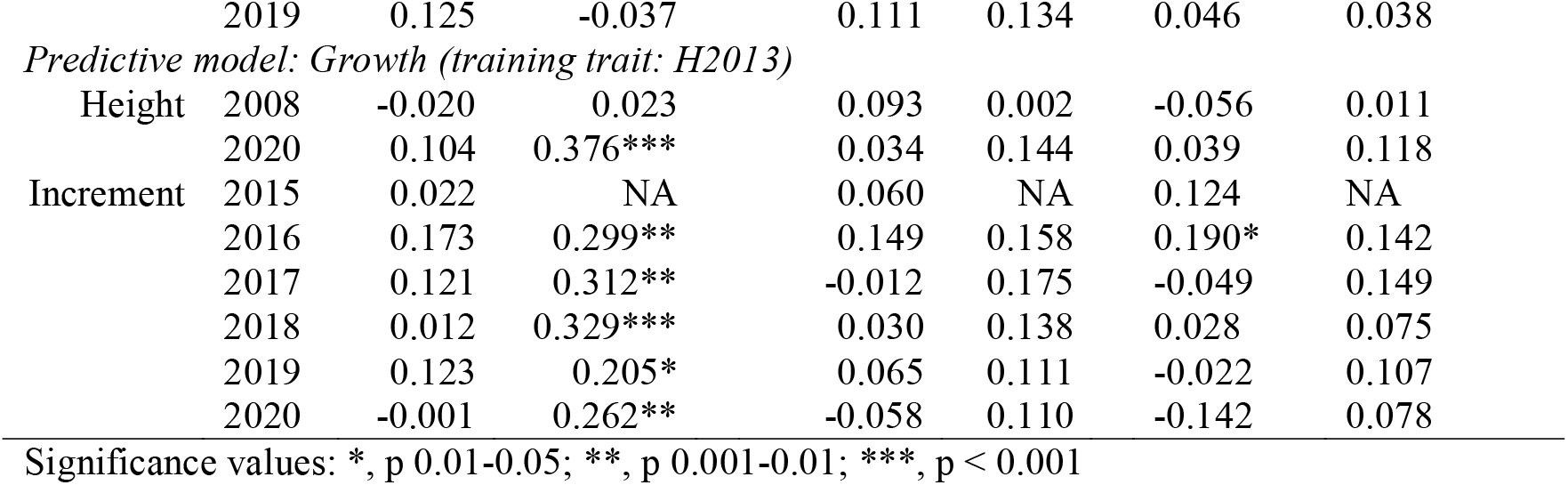
Pearson’s correlation coefficient (r) and associated significance values for comparison of predicted and observed values for each trait. Predicted values estimated by final predictive models for growth and budburst constructed using SNPs significantly associated with the traits and assessed for their performance in an internal test. Predictive models constructed using all available SNPs (no MAF filter applied, N SNPs = 15,019) trained using the full SY dataset and also trained with only SY trees from Scotland. Duration: time taken for each tree to progress from stage 4 to stage 6. Timing: time taken to reach stage 6 of budburst. Description of each budburst stage is given in Table S4.

The predicted values for growth were found to be significantly associated with observed increment measurements at YA in every year, but not at GS (Table 4). The correlation between predicted values and observed height in 2008 (at age one) was not significant at either YA or GS, despite the strong correlation observed between predicted values and observed height at age 13 at YA (Figure 4) indicating that the cumulative effect of the trees growing in the environment at YA contributed to the strength of the association.

**Figure 4.**
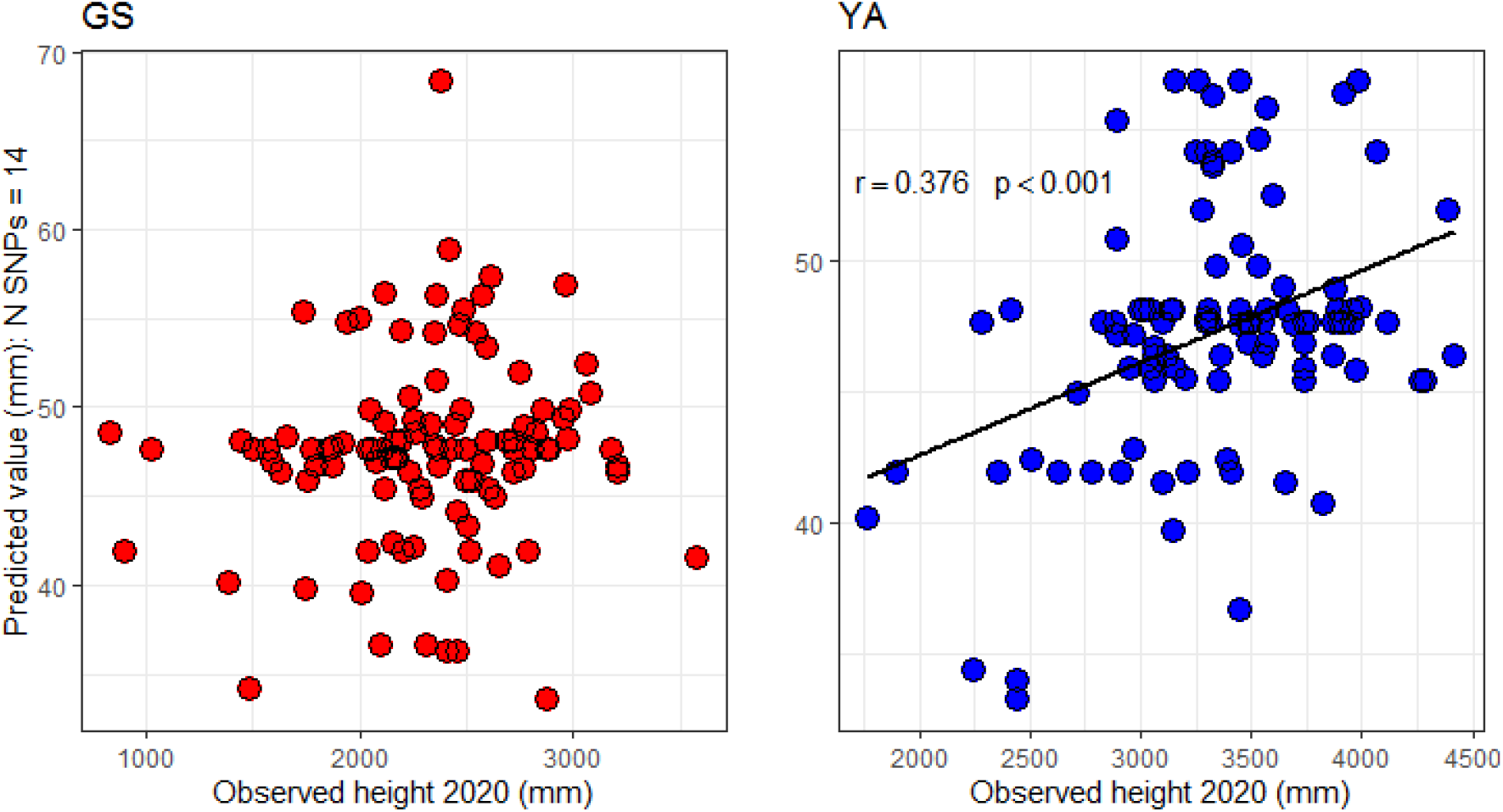
Correlations of observed height measured in 2020 at age 13 against predicted values using the final predictive model for growth for trees in an independent trial at Glensaugh (GS, correlation not significant) and Yair (YA).

The effectiveness of the final predictive model for growth as a genomic selection tool was tested by comparing different selection methods (Figure 5) in trees at both independent trial sites (GS and YA). Genomic selection was the most successful method of selecting tall trees growing at YA: trees were on average 6.80 % taller (227 mm) than trees selected using the phenotype method and 5.41 % (181 mm) taller than the mean height of trees at this site. In contrast, the phenotype method was more successful than the genomic method at GS (5.07 % and 7.70 % increase in the mean height of trees selecting using the phenotype method compared to the genomic method and site mean, respectively), despite the lack of significant correlation between height at 2008 and height at 2020 (Table S10). The coefficient of variation (CV) for trees chosen using the phenotype selection method was over 60 % greater than for those chosen using the genomic selection method at GS (23.68 and 14.63, respectively), indicating that trees chosen using the phenotype method were more variable for this trait at the site. Trees selected using the genomic and phenotype selection methods at YA had very similar CVs (10.84 and 10.31, respectively). Using the phenotype selection method, there were three trees at GS and none at YA that were among the ten tallest trees at each site. The genomic selection method identified one tree at GS and two trees at YA which were among the ten tallest trees at each site. The selected trees were from all five of the genotyped populations and included trees from 28 of the 40 available families. The majority of families were only represented by a single tree, although there were exceptions: two individuals were selected from single families in each of the sites using the phenotype method; two individuals were selected from each of two families in GS and from each of three families in YA using the genomic method.

**Figure 5.**
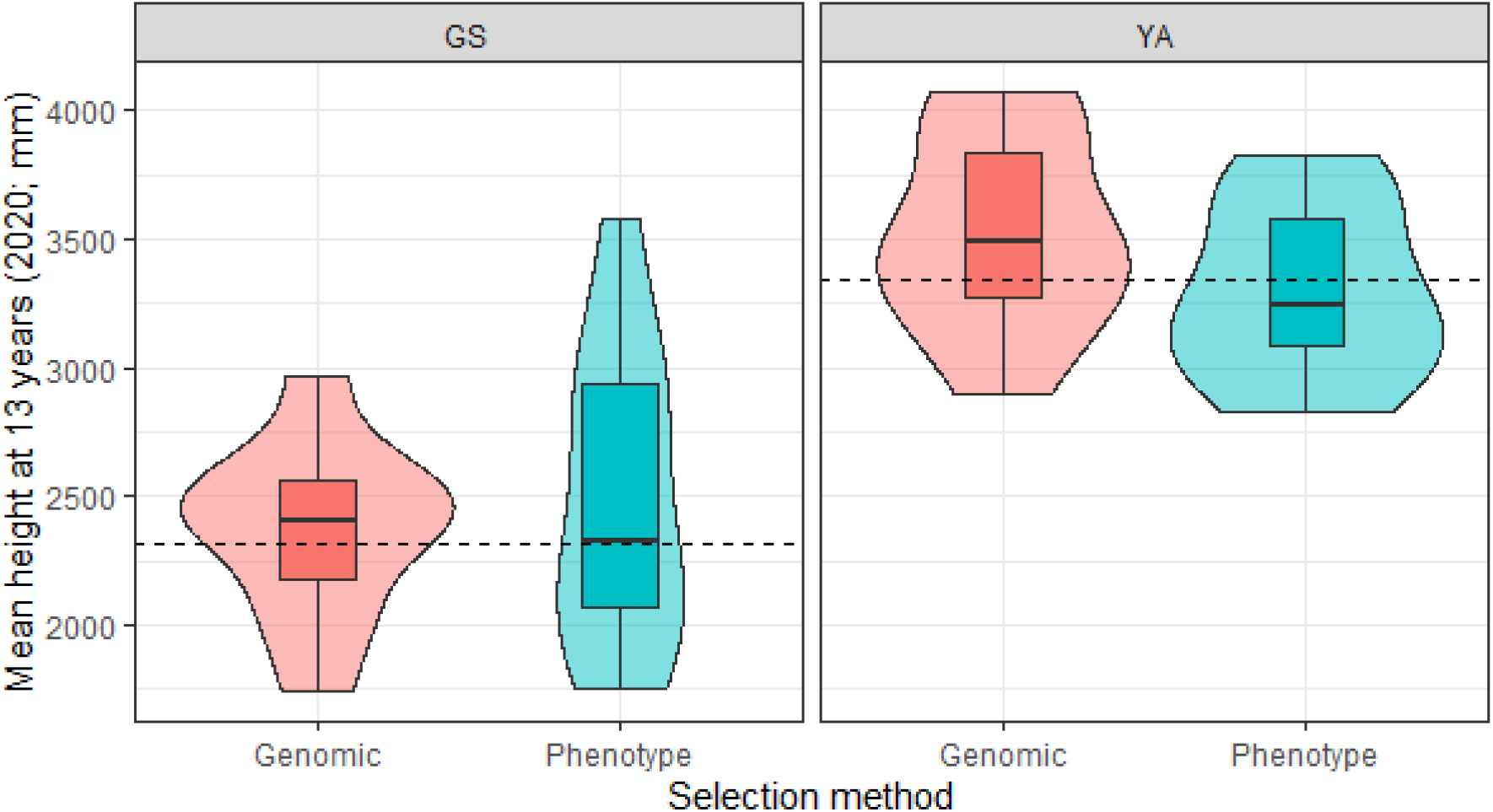
Height at 13 years (measured before the growing season started in 2020) of 10 trees at Yair (YA) and Glensaugh (GS) selected using different methods: Genomic: genomic selection to identify the predicted 10 tallest trees using values from the final predictive model for growth (SNPs identified in both SY and MU-SY-UN, no MAF filter applied, N SNPs = 14); Phenotype: phenotype selection where the 10 tallest trees at one year were selected (before the start of the second growing season, 2008). The dotted line represents the mean height of trees at each site.

## Discussion

This study is among the first to use a high throughput genotyping array to identify SNPs associated with growth and phenology traits in conifers and is unique in applying a multispecies approach. Association genetics of adaptive traits is of great interest to forestry and is being studied in many species such as *Pinus contorta* (Mahony et al 2019), *Populus trichocarpa* (Evans et al., 2014), *Picea sitchensis* (Holliday et al., 2010) and *Pinus taeda* (Lu et al., 2017) and the use of multiple species has the potential to improve the generality of models based upon them. Although other multispecies genotyping arrays have been developed (e.g. for *Eucalyptus*, Silva-Junior et al 2015), association analyses are conventionally restricted to a single species.

The high-throughput SNP array allowed nearly 50,000 SNPs (of which 20,795 were successfully converted) to be simultaneously genotyped in a large number of trees. To increase the sample size of the datasets and the statistical power of our analyses, data from *P. mugo* and *P. uncinata*, which are both part of the *P. mugo* complex, were combined. The dropout rate for *P. mugo* was much higher, and the call rate much lower than for *P. sylvestris* and *P. uncinata*. It is likely that this is a consequence of the dominance of *P. sylvestris* in the sample set used to set allele calling thresholds, coupled with the genetic distance between the two species (Perry et al., 2020). Despite this, nearly a third of SNPs on the array were high quality in all three species and nearly half of all successfully converted SNPs were polymorphic in all three species – twice the number reported by Perry et al., (2020) although sample sizes were much larger in this study.

We used the SNP datasets to test for associations with previously published phenotypes for the three pine species (Wachowiak et al., 2018a), identifying 118 SNPs significantly associated with variation in growth and phenology over multiple years, of which nearly half would not have been identified without a multispecies approach. As shown previously, the between-population variation in both phenology and height was far less in *P. mugo* and *P. uncinata* than in *P. sylvestris*, reflecting the fact that the latter was sampled from across its much broader geographical distribution and across much wider environmental gradients in photoperiod and temperature (Wachowiak et al., 2018a). Despite the smaller environmental gradient represented by our *P. mugo* and *P. uncinata* sampling, the number of SNPs identified as significantly associated with phenology was similar to the number of SNPs identified in *P. sylvestris*, although the number of SNPs identified as significantly associated with growth traits was much higher in *P. sylvestris*. The majority of studies on genetic control of adaptive traits in conifers have also identified multiple QTLs or SNPs associated with variation in timing of bud set, budburst and growth (Bartholomé et al., 2016, Eckert et al., 2009, Holliday et al., 2010, Hurme et al., 2000, Jermstad et al., 2001, Jermstad et al., 2003, Plomion et al., 1996, Prunier et al., 2013) as expected of complex traits (Mackay, 2001). For example, Eckert et al., (2015) tested 475 SNPs and found six significant associations with height and budburst in sugar pine (*Pinus lambertiana*) and Budde et al., (2014) identified 17 SNPs significantly associated with serotiny in maritime pine (*Pinus pinaster*) using an array with 251 SNPs from candidate genes. However, there have also been a limited number of specific genes implicated in the control of adaptive traits in conifers: loci related to budburst/set were identified in *Picea abies* and *Pinus sylvestris* (PaFTL2, (Avia et al., 2014) and PsFTL2, (Gyllenstrand et al., 2007), respectively).

Overall, the majority of SNPs identified in this study were rare. Of those that were common, counts were similar among *P. sylvestris* and the *P. mugo* complex for both phenology (13 and 10 for *P. sylvestris* and the *P. mugo* complex, respectively) and growth (four and six for *P. sylvestris* and the *P. mugo* complex, respectively). Although one SNP was found to be associated with both phenology and growth (the former in the *P. mugo* complex and the latter in *P. sylvestris*) it was extremely rare in *P. sylvestris*. Most likely, this is a confounding effect due to a small number of individuals (in this case, two) with a rare allele at the locus, that are at the tail-end of a trait distribution (the two individuals were ranked 366 and 412 out of 413 for increment in 2013). Although these findings seem to support the use of MAF filtering, applying a filter prior to association analyses was found to significantly reduce the number of common SNPs identified, probably as a result of changes to the PC scores and kinship matrix (describing population structure and relatedness) caused by the removal of rare variants. A further benefit of retaining all SNPs at all stages of analyses was to enable the evaluation of the relative contribution of rare and common SNPs to each trait and to assess the predictive power of models constructed using SNPs with and without MAF filtering. There were very few instances of the same SNP being associated among traits, among species or among years, a finding also reported by Westbrook et al., (2013), possibly indicating the involvement of different genes at different stages of development or in response to varying environmental conditions, as well as the very small effect sizes of most SNPs in polygenic traits (Korte and Farlow, 2013). Our earlier comparative genetic studies of a large set of SNPs located in nuclear genes similarly found almost no shared polymorphisms under selection between different taxa of the *P. mugo* complex (Wachowiak et al., 2018b).

Phenological variation in *Pinus* spp. in common garden studies has been shown to be significantly associated with the environment at the site of origin (Howe et al., 2003, Hurme et al., 1997, Repo et al., 2000, Salmela et al., 2011, Wachowiak et al., 2018a) with trees from northern European populations setting bud and flushing earlier than trees from more southerly populations. Whereas environmental cues are expected to play an important role in initiating phenological processes (Dougherty et al., 1994) including budburst (Laube et al., 2014), bud set is thought to be endogenous in *Pinus* spp, with photoperiod and temperature having relatively minor effects (Cooke et al., 2012). In this study, we found a high proportion of common SNPs in genes putatively involved in environmental responses (including response to abiotic and biotic stress and environmental cues) for both budburst and growth, but not for bud set. Common SNPs associated with bud set were exclusively located in genes related to growth and development. At this stage, assigning unigenes in conifers is largely presumptive and relies on similarity to domains or families of proteins with a large and/or speculative range of functions, many of which are, as yet, unexplored or undefined. However, the divergence of assignment among SNPs associated with budburst and bud set, and its concurrence with physiological understanding of these functions, suggests the assignment is plausible. Furthermore, as it has previously been demonstrated that intragenic linkage disequilibrium (LD) decays rapidly in our species (Wachowiak et al., 2009, Wachowiak et al., 2013), there is a higher likelihood that SNPs identified are directly involved in variation of phenology and growth. At present, our ability to better characterise the SNPs, for example determining whether they are synonymous or nonsynonymous, is limited by the paucity of highly similar, well characterised and published protein and gene sequences for these species.

Although predictive models constructed using all available polymorphic SNPs were the most successful at predicting values in the internal validation set they had no predictive ability when tested in an independent set of trees, possibly reflecting the divergent geographic ranges and associated environments of populations used in the trials (although training the models using trees from Scotland to reflect the geographic range of populations in the independent trial resulted in almost no improvement to the models’ predictive ability). In contrast, predictive models constructed using SNPs identified as significantly associated with budburst and growth in the association trial were found to be successful at estimating values in both the internal assessment and the independent assessment, although in the latter the predictive ability of the models varied spatially (among the sites) and temporally (among years). The final predictive models, chosen for their performance in the internal assessment, comprised SNPs from all species’ datasets indicating that the multispecies approach to identify SNPs was justified. When testing these models in an independent trial, observed values for height at age 13 and increment, over multiple years, were highly significantly correlated with predicted values generated by the final predictive model for growth, although only at YA. In contrast, the predictive ability of the growth model for trees at GS was poor. Phenotypic variation is a product of both heritable genetic and environmental variation. Furthermore, variation in phenotypic plasticity may cause families and populations to respond to environmental variation in different ways (Cooper et al., 2018; Gratani, 2014). Consequently, the predictive ability of models will depend on the interplay between the underlying genetic control of the traits, a host of external cues and stresses that directly and indirectly determine trait expression, and differences among the environments of trees used for association analyses and those used for external testing of predictive models. Trees growing at the YA site are much larger than at GS, indicating that there may be environmental limitations for growth at GS which are not present at YA. The trees grown in the glasshouse which were used to identify SNPs associated with growth are similarly unlikely to have experienced many environmental limitations. The lack of environmental limitations for trees growing in both the glasshouse (association trial) and at the YA site may explain why the predictive model works well in this set of trees, but doesn’t have any predictive ability when tested in trees grown under a more limiting environment at GS. Ideally, therefore, a predictive model should be used in populations from very similar environments as the population used to perform association analyses (Resende et al., 2012b). For instance, a predictive model for serotiny constructed by Budde et al., (2014) also had variable success when applied to different populations of *Pinus pinaster*. It is also possible that optimisation of the prediction models using variable selection approaches such as LASSO, would improve results, particularly where genotype x environment (G x E) interactions are likely to impact the association analyses and/or predictions (Crossa et al 2017).

Furthermore, the age of the trees used to identify SNPs associated with traits should also be considered, with respect to maternal effects which may be more significant at younger ages (Vivas et al., 2020) resulting in phenotypes which are less a product of their genotype (and environment) than in later life stages. Many more SNPs were identified as significantly associated with height and increment in 2013 than in 2011 or 2012 and an incremental reduction in the strength of the relationship between growth in the first year and in the two subsequent years was also observed: both these findings suggest that maternal effects were present in at least the first year of growth but that the effect was much less by the third year of growth. The lack of predictive ability in the final predictive model for growth in the independent trial at YA for trees in their first year of growth suggests that maternal effects may be significant in these trees, but that the effect has diminished by age 13 when the predictive ability is very good. As Lee (1999) found that height at 13 years in another commercial conifer species was a good predictor of height at final harvest the fact that our model has high predictive ability in trees at age 13 indicates that it has the potential to be a useful tool for early selection for height at final harvest in Scots pine.

The relationship between bud burst timing and duration was found to vary as budburst progressed: trees which were observed to reach the first few stages of budburst (where scales were open but needles not yet visible) early in the season did not complete the whole budburst process sooner as might be expected. Instead, these trees took longer overall to complete budburst and it is clear that this relationship is not consistent among sites, which emphasises the need for caution in applying genotype-trait relationships across environments. Similarly, the prediction model for budburst had variable accuracy among the two independent field sites: the predicted values were significantly (albeit only weakly) positively correlated with the duration of budburst for two years at GS, but not at YA, while the predicted values for budburst were significantly correlated with timing of budburst but only at YA in one year. This was a negative relationship, such that trees that were predicted to complete budburst early in the season actually completed budburst late. Although this initially seems surprising, it does have a plausible biological explanation. The predictive model was constructed using SNPs which were identified as significantly associated with the timing of budburst in a set of trees from a common garden glasshouse experiment, whilst the independent trial data were collected from trees planted outdoors in a field trial. The environmental difference between the glasshouse and the field was clearly substantial, with possibly the most important deviation between the two being that temperatures in the glasshouse did not drop below freezing throughout the winter. The relationship between the chilling requirement (the accumulation of time spent below a certain temperature) and the initiation of budburst is complex: tree species and populations differ in their chilling requirement as well as in their forcing requirement (the accumulation of time spent above a certain temperature) after the chilling requirement is met (Körner, 2006). An increase in chill days (mean temperature < 5 °C) can significantly advance budburst timing in *P. sylvestris* (Laube et al., 2014). Heritable genetic variation in the timing of budburst is therefore likely to be strongly influenced by environmental cues including chilling and subsequent forcing. The contrast between the two environments means that trees requiring a greater number of chill days before the initiation of budburst will experience a delay in the glasshouse but burst bud earlier in the field, resulting in a negative relationship among trait values in the two environments. Moreover, variation in the climate ensures that chilling and forcing conditions vary among sites as well as annually. Although the mean number of annual chill days is higher in GS than YA, GS also has fewer growing degree days which may delay the onset of budburst in some families or populations.

We found, as has been previously reported (Calleja-Rodriguez et al. 2020), that predictive ability in *P. sylvestris* (estimated as the correlation between the genomic estimated breeding values and phenotypes) was positively associated with narrow sense heritability of the trait. In contrast, the predictive ability of the models in an independent multi-site trial was not correlated with the predictive ability in the association dataset, possibly because of the different environments involved. However, the heritability estimates are extremely high for some traits (particularly bud set in *P. sylvestris*) which could be due to the distribution of SNP effect sizes (Young et al., 2018) or the average linkage disequilibrium between SNPs and causal variants being different than it is among SNPs (Evans et al., 2018). As previously noted, LD decays rapidly in these species and this may indicate that there is a higher rate of LD between SNPs and causal variants than among SNPs. Our finding that phenological traits (budburst and bud set) had higher narrow sense heritability than growth traits (height and annual increment) has also been reported in *Quercus robur* (Scotti-Saintagne et al., 2004). Similarly, high narrow sense heritability for budburst has been estimated in other conifers (*Picea abies, h*^*2*^ = 0.8: Aitken and Hannerz, 2001) as has moderate narrow sense heritability for height (*Pinus pinaster, h*^*2*^ = 0.37: Vazquez-Gonzalez et al., 2021). Variation in narrow sense heritability across years, as was observed in this study, was also reported for *Quercus robur* (*h*^*2*^ = 0.48 - 0.80; Bogdan et al., 2004) and *Pinus taeda* (*h*^*2*^ = 0 - 0.75; Balocchi et al., 1993), so we might expect the accuracy of genomic prediction to vary considerably by species and by trait.

Predictive models potentially provide a tool with which to determine the phenotype of trees at early life stages, saving both time and money. The gain of nearly 7 % in height observed using genomic selection as opposed to phenotype selection is slightly lower than the gain predicted for material derived from existing seed orchards (8-12 %: Lee, 1999) but without the extensive and expensive trial set up and maintenance. Furthermore, the height at harvest of Scots pine (with the average yield class for this species of 10) could be expected to increase by 1.08 to 1.24 m when using predictive modelling based on an average harvest height of 20 to 23 m (McLean, 2019). Predictive models have several potential applications including selecting for key traits in commercial breeding programmes and assessing forests for their response to abiotic and biotic stress. However, our results show the extent to which values generated by predictive models can vary in the strength of their correlation with the observed values depending on the environment in which they are tested. This may limit the deployment of genomic prediction across environments, but also where environment changes over time: something that will be a widespread issue in the near future (Franklin et al., 2016). Using a multispecies approach also highlights the improvements in both numbers of SNPs identified as significantly associated with SNPs and the accuracy of prediction models constructed when using SNPs from multiple species’ datasets. However, the small-scale comparisons between selection methods demonstrated the potential for predictive growth models to successfully select taller trees at one of our sites. As we had only small sample sizes and a relatively small pool of trees from which to select, the approach will require further testing using a larger set of trees in future trials.

## Conclusions

Despite its ecological and economic importance, this study is among the first to explore the association between SNPs and key adaptive traits in *P. sylvestris*, demonstrating the utility of the *Pinus* spp. high throughput array (Perry et al., 2020) for identifying genes and SNPs associated with phenology and growth traits. Development of a predictive model and validation in an independent trial furthermore demonstrates the potential of the approach for accelerated tree breeding. However, the study also highlights the limitations imposed by genotype by environment interactions. This may affect the application of predictive models in populations experiencing different environments from those in which the models were trained. Applying both a conventional single species and a novel multispecies approach to association analyses and predictive modelling exposes the constraints of the former and benefits of the latter. These results offer promise for this approach, highlighting the potential for improvement of economic traits in Scots pine and justifying future genomic studies in this species.

## Data accessibility

Phenotypes, sampling locations and SNPs have been uploaded to the EIDC (https://eidc.ac.uk/)

## Notes

https://doi.org/10.5285/55118e26-cf5c-41d6-9157-738fce6bdddf

https://doi.org/10.5285/52248442-a50f-4fc0-a73e-31c61cd27df9

### Competing Interest Statement

The authors have declared no competing interest.

### Summary of Updates

Minor changes have been made to the text to explain that traits were measured over multiple years to account for maternal effects. Additional minor changes have been made following suggestions by reviewers.

