## Supplementary materials for "Identifying and testing marker-trait associations for growth and phenology in three pine species: implications for genomic prediction"

**Table S1. Phenotypes and counts for populations (Pop), families (Fam/Pop: families per population) and total number of individuals (N) from the association trial and independent trials including all trees in the trial and those genotyped with a call rate (CR) > 80 %. Phenotypes: H, height; BB, budburst; BS, bud set.**

|  |  | All trees in trial | | | |  | | Genotyped trees CR > 80% | | |
| --- | --- | --- | --- | --- | --- | --- | --- | --- | --- | --- |
| Trial | Phenotypes | Pop | Fam/Pop | N |  | | Pop | | Fam/Pop | N |
| *Association trial* | | | | | | | | | | |
| Species: MU | H, BB, BS | 9 | 3-5 | 659 |  | | 5 | | 3-5 | 115 |
| Species: SY | H, BB, BS | 13 | 3-5 | 1031 |  | | 12 | | 3-5 | 456 |
| Species: UN | H, BB, BS | 6 | 5 | 675 |  | | 5 | | 5 | 191 |
| *Independent trial* | | | | | | | | | | |
| Site: GS | H, BB | 21 | 8 | 672 |  | | 5 | | 8 | 108 |
| Site: YA | H, BB | 21 | 8 | 672 |  | | 5 | | 7-8 | 100 |

**Table S2. Species mean and standard error for phenology (bud set: BS2010; bud burst: BB2011-2012) and growth (height: H2011-13; annual increment: I2012-13) for trees genotyped (call rate > 80 %) and analysed in this study. MU: *P. mugo*; SY: *P. sylvestris*; UN: *P. uncinata*.**

| Trait | MU | SY | UN | All species |
| --- | --- | --- | --- | --- |
| BS2010 | 19.09 ± 1.08 | 38 ± 0.76 | 27.37 ± 0.77 | 32.38 ± 0.58 |
| BB2011 | 31.64 ± 0.73 | 34.44 ± 0.36 | 37.06 ± 0.33 | 34.68 ± 0.26 |
| BB2012 | 33.56 ± 1.31 | 39.74 ± 0.78 | 48.73 ± 0.85 | 41.11 ± 0.58 |
| H2011 | 6.28 ± 0.25 | 14.08 ± 0.28 | 9.03 ± 0.22 | 11.62 ± 0.21 |
| H2012 | 16.78 ± 0.70 | 34.44 ± 0.63 | 24.43 ± 0.57 | 29.24 ± 0.48 |
| H2013 | 24.80 ± 1.03 | 48.76 ± 0.75 | 32.69 ± 0.80 | 40.89 ± 0.63 |
| I2012 | 10.49 ± 0.54 | 20.37 ± 0.43 | 15.41 ± 0.44 | 17.63 ± 0.32 |
| I2013 | 7.97 ± 0.48 | 14.31 ± 0.30 | 8.41 ± 0.33 | 11.82 ± 0.24 |

**Table S3. Average climatic variables at Glensaugh (GS) and Yair (YA) multi-site experimental field trials since planting in 2012 until 2019. Climatic variables are derived from data provided by the Met Office (daily mean, minimum and maximum temperatures and monthly rainfall).**

| Climatic variable: average 2012-19 | GS | YA |
| --- | --- | --- |
| Mean daily temperature (deg C) | 8.12 | 8.58 |
| Minimum daily temperature (deg C) | 4.85 | 4.96 |
| Maximum daily temperature (deg C) | 11.37 | 12.25 |
| Mean number of chill days (< 5 deg C) | 108.13 | 96.25 |
| Annual rainfall (mm) | 1096.68 | 962.76 |
| Growing season length (days)^1^ | 261.50 | 279.50 |
| Growing degree days^2^ | 1468.54 | 1662.84 |

^1^Growing season length: period bounded by daily mean temperature > 5 °C for > 5 consecutive days and daily mean temperature < 5 °C for > 5 consecutive days (after 1 July)

^2^Growing degree days: the mean number of degrees by which the air temperature has gone above 5 °C calculated day by day and summed over the year

**Table S4. Stages of budburst. Due to missing data, only stages 4 to 6 are analysed further**

| Stage | Description |
| --- | --- |
| 1 | Dormant |
| 2 | Bud swelling |
| 3 | Scales open at the base |
| 4 | Scales open along length of shoot, no needles |
| 5 | White tipped needles visible |
| 6 | Green needles |
| 7 | Needle separation |

**Table S5. Phenotypes and genotypes for budburst, bud set and height for all genotyped *P. sylvestris*, *P. mugo* and *P. uncinata* trees from the association trial (https://doi.org/10.5285/55118e26-cf5c-41d6-9157-738fce6bdddf)**

**Table S6. Narrow sense heritability (*h^2^*), associated standard error (SE)** **estimates for phenology and growth traits (budburst: BB; bud set: BS; height: H; annual increment: I) for years 2011-2013 for the *Pinus mugo* complex (*Pinus mugo*, MU; *Pinus uncinata*, UN), *Pinus sylvestris* (SY) and all species combined (MU-SY-UN)**

|  |  | *h^2^* ± SE | | |
| --- | --- | --- | --- | --- |
| Trait | Year | MU-UN | SY | MU-SY-UN |
| *Phenology* | | | | |
| BB | 2011 | 0.79 ± 0.15 | 0.79 ± 0.10 | 0.64 ± 0.07 |
|  | 2012 | 0.51 ± 0.14 | 0.71 ± 0.10 | 0.57 ± 0.07 |
| BS | 2010 | 0.44 ± 0.14 | 1.00 ± 0.08 | 0.91 ± 0.05 |
| *Growth* | | | | |
| H | 2011 | 0.86 ± 0.12 | 0.88 ± 0.09 | 0.57 ± 0.07 |
|  | 2012 | 0.75 ± 0.13 | 0.67 ± 0.11 | 0.49 ± 0.07 |
|  | 2013 | 0.77 ± 0.13 | 0.75 ± 0.11 | 0.58 ± 0.07 |
| I | 2012 | 0.63 ± 0.15 | 0.45 ± 0.11 | 0.34 ± 0.07 |
|  | 2013 | 0.44 ± 0.17 | 0.44 ± 0.11 | 0.32 ± 0.07 |

**Table S7. Phenotypes and genotypes for budburst and height for all genotyped *P. sylvestris* trees from the multi-site independent field trial (https://doi.org/10.5285/52248442-a50f-4fc0-a73e-31c61cd27df9)**

**Table S8. Pairwise Pearson’s correlation coefficient (r) and associated significance values for comparison of each stage of budburst and duration of budburst (time taken for each tree to progress from stage 4 to stage 6: description of each stage is given in Table 1) for each year from 2015 to 2019 at each site**

|  |  | GS |  |  |  | YA |  |  |
| --- | --- | --- | --- | --- | --- | --- | --- | --- |
| Year | Stage | Duration | Stage 4 | Stage 5 |  | Duration | Stage 4 | Stage 5 |
| 2015 | 4 | -0.789*** |  |  |  | -0.823*** |  |  |
|  | 5 | -0.213* | 0.567*** |  |  | -0.161 | 0.662*** |  |
|  | 6 | 0.348*** | 0.302** | 0.514*** |  | 0.068 | 0.510*** | 0.919*** |
| 2016 | 4 | 0.091 |  |  |  | -0.928*** |  |  |
|  | 5 | 0.445*** | 0.492*** |  |  | -0.557*** | 0.708*** |  |
|  | 6 | 0.937*** | 0.432*** | 0.585*** |  | -0.065 | 0.431*** | 0.548*** |
| 2017 | 4 | -0.354*** |  |  |  | -0.927*** |  |  |
|  | 5 | 0.043 | 0.524*** |  |  | -0.429*** | 0.627*** |  |
|  | 6 | 0.609*** | 0.526*** | 0.521*** |  | 0.071 | 0.309** | 0.474*** |
| 2018 | 4 | -0.549*** |  |  |  | -0.569*** |  |  |
|  | 5 | -0.070 | 0.675*** |  |  | 0.055 | 0.615*** |  |
|  | 6 | 0.382*** | 0.563*** | 0.677*** |  | 0.529*** | 0.394*** | 0.701*** |
| 2019 | 4 | 0.021 |  |  |  | -0.667*** |  |  |
|  | 5 | 0.479*** | 0.535*** |  |  | -0.371*** | 0.793*** |  |
|  | 6 | 0.886*** | 0.483*** | 0.668*** |  | 0.363*** | 0.451*** | 0.558*** |

Significance values: *, p 0.01-0.05; **, p 0.001-0.01; ***, p < 0.001. Site codes: GS, Glensaugh; YA, Yair. Description of each stage of budburst is given in Table 1.

**Table S9. Pearson’s correlation coefficient (r) and associated significance (*, p 0.01-0.05; **, p 0.001-0.01; ***, p < 0.001) for growth and phenology traits measured in genotyped trees (CR > 80 %) for the glasshouse trial for each species separately (MU, *Pinus mugo*; SY, *Pinus sylvestris*; UN, *Pinus uncinata*). Traits: BS, bud set; BB, budburst; H, height; I, annual increment.**

| Species | Trait | BS2010 | BB2011 | BB2012 | H2011 | H2012 | H2013 | I2012 |
| --- | --- | --- | --- | --- | --- | --- | --- | --- |
| MU | BB2011 | -0.34*** |  |  |  |  |  |  |
|  | BB2012 | 0.10 | 0.30** |  |  |  |  |  |
|  | H2011 | -0.10 | 0.15 | -0.07 |  |  |  |  |
|  | H2012 | 0.04 | 0.13 | -0.06 | 0.74*** |  |  |  |
|  | H2013 | 0.04 | 0.09 | -0.11 | 0.67*** | 0.91*** |  |  |
|  | I2012 | 0.10 | 0.09 | -0.04 | 0.49*** | 0.95*** | 0.87*** |  |
|  | I2013 | -0.01 | 0.01 | -0.10 | 0.36*** | 0.49*** | 0.80*** | 0.47*** |
| SY | BB2011 | 0.28*** |  |  |  |  |  |  |
|  | BB2012 | 0.25*** | 0.42*** |  |  |  |  |  |
|  | H2011 | 0.17*** | 0.29*** | 0.32*** |  |  |  |  |
|  | H2012 | 0.21*** | 0.15** | 0.16*** | 0.81*** |  |  |  |
|  | H2013 | 0.18*** | 0.10* | 0.08 | 0.73*** | 0.91*** |  |  |
|  | I2012 | 0.19*** | 0.03 | 0.03 | 0.53*** | 0.92*** | 0.85*** |  |
|  | I2013 | -0.05 | -0.09 | -0.17*** | 0.04 | 0.09 | 0.48*** | 0.10* |
| UN | BB2011 | -0.46*** |  |  |  |  |  |  |
|  | BB2012 | -0.04 | 0.27*** |  |  |  |  |  |
|  | H2011 | -0.02 | 0.09 | 0.10 |  |  |  |  |
|  | H2012 | 0.00 | -0.04 | -0.10 | 0.71*** |  |  |  |
|  | H2013 | -0.04 | 0.00 | -0.14 | 0.66*** | 0.93*** |  |  |
|  | I2012 | 0.00 | -0.09 | -0.18* | 0.41*** | 0.94*** | 0.88*** |  |
|  | I2013 | -0.06 | 0.02 | -0.15* | 0.32*** | 0.48*** | 0.77*** | 0.47*** |

| **Table S10. Peason’s correlation coefficient (r) and associated significance (*, p 0.01-0.05; **, p 0.001-0.01; ***, p < 0.001) for growth and phenology traits over multiple years measured in genotyped trees for the independent multi-site field trial for each site separately (GS, Glensaugh - values below the diagonal; YA, Yair - values above the diagonal). Traits: H, height; I, annual increment; T, timing of budburst, time to reach stage 6; D, duration of budburst: time to progress from stage 4 to 6 (see Table S4 for definitions of each budburst stage).** | | | | | | | | | | | | | | | | | | |
| --- | --- | --- | --- | --- | --- | --- | --- | --- | --- | --- | --- | --- | --- | --- | --- | --- | --- | --- |
| Trait | H08 | H20 | I15 | I16 | I17 | I18 | I19 | I20 | T15 | T16 | T17 | T18 | T19 | D15 | D16 | D17 | D18 | D19 |
| H08 |  | 0.17 | NA | 0.15 | 0.13 | 0.09 | 0.06 | 0.07 | 0.11 | 0.20 | 0.01 | -0.10 | 0.06 | 0.06 | -0.01 | -0.02 | 0.03 | 0.03 |
| H20 | 0.12 |  | NA | 0.76  *** | 0.84  *** | 0.82  *** | 0.76  *** | 0.67  *** | 0.03 | -0.35  *** | 0.01 | -0.01 | -0.14 | 0.11 | -0.12 | 0.12 | 0.08 | -0.14 |
| I15 | 0.09 | 0.64  *** |  | NA | NA | NA | NA | NA | NA | NA | NA | NA | NA | NA | NA | NA | NA | NA |
| I16 | 0.02 | 0.70  *** | 0.55  *** |  | 0.75  *** | 0.59  *** | 0.44  *** | 0.34  *** | 0.08 | -0.43  *** | -0.11 | 0.04 | -0.14 | 0.13 | -0.06 | -0.01 | 0.11 | -0.06 |
| I17 | 0.23  * | 0.71  *** | 0.46  *** | 0.53  *** |  | 0.70  *** | 0.56  *** | 0.44  *** | -0.10 | -0.38  *** | -0.08 | -0.10 | -0.21  * | 0.14 | 0.06 | 0.12 | 0.07 | -0.13 |
| I18 | 0.06 | 0.74  *** | 0.24  * | 0.52  *** | 0.48  *** |  | 0.63  *** | 0.47  *** | -0.09 | -0.34  *** | 0.16 | -0.08 | -0.09 | 0.11 | -0.04 | 0.22  * | 0.01 | -0.08 |
| I19 | 0.01 | 0.76  *** | 0.33  *** | 0.45  *** | 0.40  *** | 0.49  *** |  | 0.39  *** | 0.01 | -0.20  * | -0.03 | -0.10 | -0.05 | 0.22  * | -0.06 | 0.24  * | 0.03 | -0.07 |
| I20 | 0.15 | 0.69  *** | 0.26  ** | 0.26  ** | 0.33  *** | 0.48  *** | 0.56  *** |  | 0.00 | -0.23  * | -0.04 | 0.07 | -0.21  * | 0.07 | -0.09 | 0.14 | 0.13 | -0.27  ** |
| T15 | 0.03 | -0.06 | 0.02 | -0.05 | -0.07 | -0.06 | -0.10 | -0.04 |  | 0.10 | 0.09 | 0.38  *** | 0.37  *** | 0.07 | -0.40  *** | -0.28  ** | 0.06 | -0.02 |
| T16 | 0.17 | 0.28  ** | 0.38  *** | 0.21  * | 0.20 | 0.15 | 0.07 | 0.08 | 0.42  *** |  | 0.18 | 0.11 | 0.34  *** | -0.25  * | -0.07 | -0.16 | -0.04 | 0.00 |
| T17 | 0.05 | 0.12 | 0.20  * | 0.04 | 0.16 | 0.10 | 0.11 | -0.03 | 0.35  *** | 0.66  *** |  | 0.19 | 0.16 | -0.40  *** | -0.22 | 0.07 | 0.02 | -0.01 |
| T18 | 0.14 | 0.09 | 0.20  * | 0.12 | 0.27  ** | 0.08 | -0.05 | -0.09 | 0.37  *** | 0.49  *** | 0.66  *** |  | 0.21  ** | -0.13 | -0.38  *** | -0.25  * | 0.53  *** | -0.13 |
| T19 | 0.13 | 0.19 | 0.25  ** | 0.08 | 0.12 | 0.10 | 0.08 | 0.07 | 0.26  ** | 0.59  *** | 0.55  *** | 0.42  *** |  | -0.13 | -0.34  *** | -0.26  * | -0.08 | 0.36  *** |
| D15 | -0.04 | -0.16 | -0.12 | -0.08 | -0.15 | -0.17 | -0.03 | -0.01 | 0.35  *** | -0.17 | -0.24  * | -0.20  * | -0.20 |  | 0.43  *** | 0.30  ** | 0.17 | 0.28  ** |
| D16 | 0.15 | 0.21  * | 0.35  *** | 0.19 | 0.13 | 0.07 | 0.04 | 0.04 | 0.40  *** | 0.94  *** | 0.61  *** | 0.43  *** | 0.45 | -0.09 |  | 0.41  *** | 0.05 | 0.19 |
| D17 | 0.02 | 0.09 | 0.15 | 0.10 | 0.13 | 0.03 | 0.10 | -0.01 | 0.17 | 0.32  ** | 0.61  *** | 0.29  ** | 0.12 | 0.19 | 0.37  *** |  | 0.15 | 0.09 |
| D18 | 0.17 | 0.07 | 0.10 | 0.02 | 0.16 | -0.04 | 0.12 | 0.04 | 0.17 | 0.00 | 0.15 | 0.38  *** | -0.14 | 0.28  ** | 0.01 | 0.39  *** |  | -0.21  * |
| D19 | 0.04 | 0.11 | 0.19  * | 0.05 | 0.05 | 0.05 | 0.00 | 0.04 | 0.26  ** | 0.45  *** | 0.46  *** | 0.35  *** | 0.89  *** | -0.11 | 0.35  *** | 0.14 | -0.12 |  |

**Table S11. Adjusted mean sum of squares from ANOVA for traits in the independent multi-site experimental trial at Glensaugh (GS) and Yair (YA) from 2008 to 2020 (08-20). T: budburst time to reach stage 6; D: Budburst duration to progress from stage 4 to 6; H: height; I: increment. Degrees of freedom (df) for GS: block = 3; fam(pop) = 148; pop = 20. Df for YA: block = 3; fam(pop) = 147; pop = 20. Degrees of freedom for residuals in parentheses. Associated significance: *, p 0.01-0.05; **, p 0.001-0.01; ***, p < 0.001.**

|  | Adjusted mean sum of squares for each source of variation | | | | | | | | |
| --- | --- | --- | --- | --- | --- | --- | --- | --- | --- |
|  | GS |  |  |  |  | YA |  |  |  |
| Trait | Block | Fam(Pop) | Pop | Residual |  | Block | Fam(Pop) | Pop | Residual |
| T15 | 5.153 | 7.35 | 7.50 | 6.08  (484) |  | 516.26  *** | 16.99 | 14.51 | 13.91  (482) |
| T16 | 74.39 | 51.36  ** | 76.11 | 37.23  (457) |  | 10.24 | 5.94  * | 6.07 | 4.65  (480) |
| T17 | 214.99  *** | 48.39  ** | 63.94 | 33.34  (490) |  | 24.54  ** | 4.44 | 3.83 | 4.39  (470) |
| T18 | 18.82 | 27.34  *** | 41.88 | 16.94  (489) |  | 184.01  *** | 13.21  * | 20.84 | 10.66  (482) |
| T19 | 63.48 | 54.89  *** | 90.62  * | 28.44  (485) |  | 34.83 | 20.35  * | 16.47 | 15.6  (480) |
| D15 | 21.00 | 25.96  ** | 23.49 | 18.52  (482) |  | 617.57  *** | 44.69  ** | 45.50 | 31.85  (482) |
| D16 | 26.67 | 37.83 | 63.41  * | 31.71  (457) |  | 101.80  * | 40.70  ** | 27.11 | 28.72  (480) |
| D17 | 205.35  *** | 50.43  ** | 82.51 | 33.9  (490) |  | 9.62 | 24.63 | 27.06 | 20.62  (466) |
| D18 | 19.05 | 26.82  ** | 48.23  * | 17.73  (489) |  | 176.57  *** | 12.02 | 21.32  * | 14.41  (482) |
| D19 | 68.07 | 46.58  *** | 66.18 | 26.14  (485) |  | 293.07  *** | 25.98 | 23.36 | 21.51  (480) |
| H08 | 85.65 | 207.30  *** | 735.15  *** | 102.98  (499) |  | 105.50 | 188.30  *** | 758.30  *** | 104.20  (394) |
| H20 | 305138 | 230864  * | 1175687  *** | 179298  (482) |  | 880512  * | 412877  *** | 1006598  ** | 247731  (482) |
| I15 | 4462 | 4571  * | 16003  *** | 3387  (493) |  | NA | NA | NA | NA |
| I16 | 5826 | 6350 | 19583  *** | 5140  (494) |  | 51247  * | 10689 | 16970 | 9227  (481) |
| I17 | 2325 | 7425 | 23514  *** | 7075  (492) |  | 54595  *** | 13010  ** | 14680 | 8962  (482) |
| I18 | 8722 | 10779 | 22148  ** | 9261  (488) |  | 24012 | 13969 | 32810  ** | 11389  (482) |
| I19 | 15467 | 12786  * | 42575  *** | 10058  (486) |  | 209966  *** | 23057  ** | 53843  ** | 15398  (482) |
| I20 | 44375  * | 13435 | 26766  * | 12875  (478) |  | 52957  * | 17229 | 38233  ** | 14564  (482) |

**Table S12. Statistical power of each SNP dataset.**

|  | N SNPs | N samples | Statistical power | | | | |
| --- | --- | --- | --- | --- | --- | --- | --- |
| Species |  |  | λ = 0.1 | λ = 0.5 | λ = 1 | λ = 5 | λ = 10 |
| MU | 11,272 | 115 | 0.02 | 0.02 | 0.03 | 0.07 | 0.08 |
| SY | 15,019 | 456 | 0.61 | 0.67 | 0.72 | 0.86 | 0.90 |
| UN | 15,210 | 191 | 0.07 | 0.09 | 0.12 | 0.21 | 0.26 |
| MU-UN | 16,452 | 306 | 0.27 | 0.34 | 0.41 | 0.64 | 0.71 |
| MU-SY-UN | 17,212 | 762 | 0.96 | 0.98 | 0.99 | 1.00 | 1.00 |
| SY-[MU] | 13,047 | 115 | 0.02 | 0.02 | 0.02 | 0.03 | 0.03 |
| SY-[UN] | 13,867 | 191 | 0.07 | 0.08 | 0.10 | 0.14 | 0.16 |

Species codes: MU, *P. mugo*; SY, *P. sylvestris*; UN, *P. uncinata*. Where species were combined into a single dataset these are indicated (MU-UN and MU-SY-UN). Statistical power in a reduced sample size of *P. sylvestris* was also performed to compare with equivalent sample sizes in MU (SY-[MU]) and UN (SY-[UN]). Λ = polygenic effect

**Table S13. SNPs identified using mixed linear model (MLM) and multi-locus mixed model (MLMM) approaches as significantly associated with traits (BB2011: budburst 2011; BS2010: bud set 2010; H2011: height 2011; H2012: height 2012; H2013: height 2013; I2012: increment 2012; I2013: increment 2013). SNP labels as per Perry et al., (2020). Species: MU, *Pinus mugo*; SY, *Pinus sylvestris*; UN, *Pinus uncinata*. Minor allele frequency (MAF) <0.05: rare; >0.05: common.**

| SNP | Species | MAF | Method |
| --- | --- | --- | --- |
| *Phenology: BB2011* |  |  |  |
| comp16733_c0_seq1_222 | MU-UN | Rare | MLM |
| comp18631_c0_seq1_563 | MU-UN | Common | MLM |
| comp19208_c0_seq1_2090 | SY | Rare | MLM |
| comp19471_c0_seq1_746 | MU-UN | Rare | MLM |
|  | MU-SY-UN | Rare | MLM |
| comp19484_c0_seq1_274 | SY | Common | MLM |
| comp26730_c0_seq1_2554 | SY | Rare | MLM |
| comp29799_c0_seq1_84 | MU-UN | Common | MLM |
| comp31899_c0_seq1_3609 | MU-UN | Rare | MLM+MLMM |
|  | MU-SY-UN | Rare | MLM+MLMM |
| comp37834_c0_seq1_118 | MU-UN | Rare | MLM |
|  | MU-SY-UN | Rare | MLM |
| comp38055_c0_seq1_129 | MU-UN | Rare | MLM |
| comp39493_c0_seq1_935 | MU-UN | Common | MLM |
| comp39509_c0_seq1_5557 | MU-UN | Rare | MLM |
| comp40966_c0_seq1_403 | MU-UN | Rare | MLM |
| comp41446_c0_seq1_714 | SY | Common | MLM |
| comp42063_c0_seq1_922 | MU-UN | Rare | MLM |
|  | MU-SY-UN | Rare | MLM |
| comp42130_c0_seq1_1318 | SY | Common | MLM |
|  | MU-SY-UN | Common | MLM |
| comp42558_c0_seq1_515 | MU-SY-UN | Common | MLM |
| comp42661_c0_seq1_613 | SY | Common | MLM |
| comp43117_c0_seq1_203 | MU-UN | Common | MLM |
| comp45269_c0_seq1_760 | SY | Rare | MLM+MLMM |
| comp45607_c0_seq1_1055 | SY | Rare | MLM |
| comp46049_c0_seq1_1689 | SY | Common | MLM |
|  | MU-SY-UN | Common | MLM |
| comp46858_c0_seq1_1024 | MU-UN | Common | MLM |
|  | MU-SY-UN | Rare | MLM |
| comp47733_c0_seq1_4708 | SY | Common | MLM |
|  | MU-SY-UN | Common | MLM |
| comp47733_c0_seq1_5033 | SY | Common | MLM |
| comp47733_c0_seq1_5101 | SY | Common | MLM |
| comp48008_c0_seq1_836 | MU-UN | Common | MLM |
| comp48202_c0_seq1_307 | SY | Rare | MLM |
|  | MU-SY-UN | Rare | MLM |
| comp48223_c0_seq1_312 | MU-UN | Rare | MLM+MLMM |
| comp48223_c0_seq1_5041 | MU-UN | Rare | MLM |
|  | MU-SY-UN | Rare | MLM |
| comp48223_c0_seq1_688 | MU-UN | Rare | MLM |
|  | MU-SY-UN | Rare | MLM |
| comp48335_c0_seq3_629 | MU-UN | Rare | MLM |
| comp48745_c0_seq3_659 | MU-UN | Rare | MLM |
|  | MU-SY-UN | Rare | MLM |
| comp49205_c0_seq1_2527 | MU-UN | Rare | MLM |
|  | MU-SY-UN | Rare | MLM |
| comp49247_c0_seq1_1006 | MU-UN | Rare | MLM |
| comp49635_c0_seq1_343 | SY | Rare | MLM+MLMM |
| comp50525_c0_seq1_2688 | MU-UN | Rare | MLM |
| comp51128_c0_seq1_1529 | MU-UN | Common | MLM |
|  | MU-SY-UN | Common | MLM |
| comp51290_c0_seq5_216 | MU-UN | Common | MLM |
|  | MU-SY-UN | Rare | MLM |
| comp51722_c0_seq1_418 | MU-UN | Rare | MLM |
| comp51726_c0_seq1_4360 | MU-UN | Common | MLM |
| comp52159_c0_seq1_436 | MU-UN | Common | MLM |
| comp52277_c0_seq1_1396 | MU-UN | Rare | MLM |
|  | MU-SY-UN | Rare | MLM |
| comp52760_c0_seq1_196 | MU-UN | Rare | MLM |
|  | MU-SY-UN | Rare | MLM |
| comp52794_c0_seq1_2165 | SY | Rare | MLM+MLMM |
| comp53226_c0_seq1_1405 | SY | Rare | MLM |
|  | MU-SY-UN | Rare | MLM+MLMM |
| comp53414_c0_seq1_1227 | MU-UN | Rare | MLM |
|  | MU-SY-UN | Rare | MLM |
| comp53528_c0_seq1_2969 | MU-UN | Rare | MLMM |
| comp53584_c0_seq1_2201 | SY | Common | MLM |
| comp53736_c0_seq1_314 | MU-UN | Rare | MLM |
|  | MU-SY-UN | Rare | MLM |
| comp54202_c0_seq2_791 | MU-UN | Rare | MLM |
| comp54389_c0_seq1_1101 | MU-UN | Rare | MLM |
|  | MU-SY-UN | Rare | MLM |
| comp55037_c0_seq11_249 | MU-UN | Rare | MLMM |
|  | MU-SY-UN | Rare | MLMM |
| comp55130_c0_seq108_549 | SY | Rare | MLM |
|  | MU-SY-UN | Rare | MLM |
| comp55695_c0_seq3_1742 | MU-UN | Rare | MLM |
| comp55731_c0_seq1_2351 | SY | Rare | MLM |
| comp58475_c0_seq1_843 | MU-UN | Rare | MLM |
| Doth_comp54682_c0_seq1_159_1118 | SY | Rare | MLM |
|  | MU-SY-UN | Rare | MLM |
| *BS2010* |  |  |  |
| comp18509_c0_seq1_1470 | SY | Rare | MLM |
| comp19815_c0_seq1_115 | SY | Rare | MLM |
| comp20404_c0_seq1_397 | SY | Rare | MLM |
| comp22360_c0_seq1_1093 | MU-SY-UN | Common | MLM |
| comp33699_c0_seq1_1791 | SY | Common | MLM |
| comp38529_c0_seq1_937 | SY | Rare | MLM |
| comp42043_c0_seq1_534 | SY | Rare | MLM |
| comp44180_c0_seq1_322 | SY | Rare | MLM |
| comp44449_c0_seq1_63 | SY | Rare | MLM |
| comp45158_c0_seq1_1173 | SY | Common | MLM |
| comp46712_c0_seq1_269 | SY | Rare | MLM |
| comp47469_c0_seq1_2446 | SY | Common | MLM |
| comp48583_c0_seq1_703 | SY | Rare | MLM |
| comp49104_c0_seq1_528 | SY | Rare | MLM |
| comp50197_c0_seq1_652 | SY | Rare | MLM |
| comp52528_c0_seq1_927 | SY | Common | MLM |
| comp53726_c0_seq1_54 | SY | Rare | MLM |
| comp54134_c0_seq1_728 | SY | Rare | MLM |
| comp55659_c0_seq1_1054 | SY | Rare | MLM |
| *Growth: H2011* |  |  |  |
| comp43635_c0_seq1_122 | SY | Rare | MLM |
| comp47901_c0_seq1_977 | MU-SY-UN | Common | MLM |
| comp57875_c0_seq1_2681 | MU-SY-UN | Rare | MLM |
| *H2012* |  |  |  |
| comp44927_c0_seq1_1219 | MU-SY-UN | Common | MLM |
| *H2013* |  |  |  |
| comp46813_c0_seq1_315 | MU-SY-UN | Common | MLMM |
| comp51669_c0_seq5_124 | MU-SY-UN | Common | MLM |
| comp53296_c0_seq2_1323 | MU-SY-UN | Common | MLM |
| comp53606_c0_seq1_1256 | SY | Common | MLM |
| comp57469_c0_seq1_1389 | SY | Common | MLM |
|  | MU-SY-UN | Common | MLM |
| comp78883_c0_seq1_932 | MU-SY-UN | Common | MLM |
| *I2012* |  |  |  |
| comp44927_c0_seq1_1219 | MU-SY-UN | Common | MLM |
| *I2013* |  |  |  |
| comp19094_c0_seq1_1018 | SY | Rare | MLM |
| comp19526_c0_seq1_837 | SY | Rare | MLM |
| comp19814_c0_seq1_714 | SY | Rare | MLM |
| comp20305_c0_seq1_732 | SY | Rare | MLM |
| comp31134_c0_seq1_289 | SY | Rare | MLM+MLMM |
| comp33384_c0_seq1_1437 | SY | Rare | MLM |
| comp38562_c0_seq1_1084 | SY | Rare | MLM |
| comp39306_c0_seq1_355 | MU-UN | Rare | MLM+MLMM |
|  | MU-SY-UN | Rare | MLM |
| comp40846_c0_seq1_2337 | MU-UN | Common | MLM |
|  | MU-SY-UN | Common | MLM |
| comp41840_c0_seq1_158 | SY | Common | MLM |
|  | MU-SY-UN | Common | MLM |
| comp42123_c0_seq1_2358 | MU-UN | Common | MLM |
|  | MU-SY-UN | Rare | MLM |
| comp44962_c0_seq1_395 | SY | Rare | MLMM |
| comp45287_c1_seq1_139 | MU-UN | Common | MLM |
|  | MU-SY-UN | Common | MLM |
| comp45357_c0_seq1_346 | MU-SY-UN | Rare | MLMM |
| comp45673_c0_seq1_2716 | SY | Rare | MLM |
| comp46058_c0_seq1_1011 | SY | Rare | MLM |
| comp46438_c0_seq1_1282 | SY | Rare | MLM |
| comp46813_c0_seq1_315 | MU-SY-UN | Common | MLMM |
| comp47138_c0_seq1_290 | SY | Rare | MLM |
| comp47383_c0_seq1_174 | MU-SY-UN | Common | MLM |
| comp47519_c0_seq1_2584 | SY | Rare | MLM+MLMM |
| comp49053_c0_seq1_1206 | MU-UN | Common | MLM |
|  | MU-SY-UN | Rare | MLM |
| comp49307_c1_seq1_140 | SY | Rare | MLM |
| comp50406_c0_seq1_2537 | MU-UN | Common | MLM |
| comp50686_c0_seq1_570 | SY | Common | MLM |
| comp50796_c0_seq1_1297 | SY | Rare | MLM |
| comp51128_c0_seq1_1529 | SY | Rare | MLM |
| comp51215_c0_seq1_1350 | SY | Rare | MLM |
| comp51704_c0_seq1_1790 | SY | Rare | MLM |
| comp51904_c0_seq1_501 | MU-UN | Common | MLM |
|  | MU-SY-UN | Common | MLM |
| comp51948_c0_seq1_1379 | SY | Rare | MLM+MLMM |
| comp54416_c0_seq10_954 | SY | Rare | MLM |
| comp55723_c1_seq1_3800 | SY | Rare | MLM |
|  | MU-SY-UN | Rare | MLM |
| comp78883_c0_seq1_932 | MU-SY-UN | Common | MLM |

**Table S14. Genotype frequencies for SNPs identified as significantly associated with adaptive traits in the *P. mugo* complex (*P. mugo*, MU; *P. uncinata*, UN). Genotypic class frequencies for each species are given as the proportion of calls which were homozygous (AA or BB) or heterozygous (AB) not including calls which had failed.**

|  | UN |  |  |  | MU |  |  |
| --- | --- | --- | --- | --- | --- | --- | --- |
| SNP | AA | AB | BB |  | AA | AB | BB |
| *Trait: BB2011* |  |  |  |  |  |  |  |
| comp16733_c0_seq1_222 | 0.98 | 0.02 |  |  | 0.84 | 0.16 |  |
| comp18631_c0_seq1_563 | 0.97 | 0.03 |  |  | 0.44 | 0.49 | 0.07 |
| comp19471_c0_seq1_746 | 1.00 |  |  |  | 0.75 | 0.24 | 0.01 |
| comp29799_c0_seq1_84 | 0.98 | 0.02 |  |  | 0.44 | 0.53 | 0.03 |
| comp31899_c0_seq1_3609 | 1.00 |  |  |  | 0.98 | 0.02 |  |
| comp37834_c0_seq1_118 | 1.00 |  |  |  | 0.99 |  | 0.01 |
| comp38055_c0_seq1_129 | 0.96 | 0.04 |  |  | 0.92 | 0.07 | 0.01 |
| comp39493_c0_seq1_935 | 0.98 | 0.02 |  |  | 0.53 | 0.43 | 0.04 |
| comp39509_c0_seq1_5557 | 1.00 |  |  |  | 0.94 | 0.06 |  |
| comp40966_c0_seq1_403 | 1.00 |  |  |  | 0.99 | 0.01 |  |
| comp42063_c0_seq1_922 | 1.00 |  |  |  | 0.99 | 0.01 |  |
| comp43117_c0_seq1_203 | 0.97 | 0.03 |  |  | 0.63 | 0.36 | 0.01 |
| comp46858_c0_seq1_1024 | 1.00 |  |  |  | 0.43 | 0.56 | 0.01 |
| comp48008_c0_seq1_836 | 0.88 | 0.12 |  |  | 0.91 | 0.08 | 0.01 |
| comp48223_c0_seq1_312 | 1.00 |  |  |  | 0.99 | 0.01 |  |
| comp48223_c0_seq1_5041 | 1.00 |  |  |  | 0.96 | 0.02 | 0.01 |
| comp48223_c0_seq1_688 | 1.00 |  |  |  | 0.77 | 0.23 | 0.01 |
| comp48335_c0_seq3_629 | 0.98 | 0.02 |  |  | 0.95 | 0.05 |  |
| comp48745_c0_seq3_659 | 1.00 |  |  |  | 0.99 | 0.01 |  |
| comp49205_c0_seq1_2527 | 1.00 |  |  |  | 0.99 | 0.01 |  |
| comp49247_c0_seq1_1006 | 0.98 | 0.02 |  |  | 0.99 | 0.01 |  |
| comp50525_c0_seq1_2688 | 1.00 |  |  |  | 0.99 | 0.01 |  |
| comp51128_c0_seq1_1529 | 0.75 | 0.25 |  |  | 0.55 | 0.44 | 0.01 |
| comp51290_c0_seq5_216 | 0.95 | 0.05 |  |  | 0.65 | 0.32 | 0.03 |
| comp51722_c0_seq1_418 | 1.00 |  |  |  | 0.99 | 0.01 |  |
| comp51726_c0_seq1_4360 | 0.92 | 0.08 |  |  | 0.69 | 0.30 | 0.01 |
| comp52159_c0_seq1_436 | 0.94 | 0.06 |  |  | 0.85 | 0.14 | 0.01 |
| comp52277_c0_seq1_1396 | 1.00 |  |  |  | 0.97 | 0.03 |  |
| comp52760_c0_seq1_196 | 1.00 |  |  |  | 0.99 | 0.01 |  |
| comp53414_c0_seq1_1227 | 1.00 |  |  |  | 0.99 | 0.01 |  |
| comp53528_c0_seq1_2969 | 1.00 |  |  |  | 0.82 | 0.18 |  |
| comp53736_c0_seq1_314 | 1.00 |  |  |  | 0.98 |  | 0.02 |
| comp54202_c0_seq2_791 | 1.00 |  |  |  | 0.99 | 0.01 |  |
| comp54389_c0_seq1_1101 | 1.00 |  |  |  | 0.99 | 0.01 |  |
| comp55037_c0_seq11_249 | 1.00 |  |  |  | 0.99 | 0.01 |  |
| comp55695_c0_seq3_1742 | 1.00 |  |  |  | 0.99 | 0.01 |  |
| comp58475_c0_seq1_843 | 0.99 | 0.01 |  |  | 0.95 | 0.05 |  |
| *Trait: I2013* |  |  |  |  |  |  |  |
| comp39306_c0_seq1_355 | 1.00 |  |  |  | 0.96 | 0.04 |  |
| comp40846_c0_seq1_2337 | 0.79 | 0.21 |  |  | 0.34 | 0.65 | 0.01 |
| comp42123_c0_seq1_2358 | 0.93 | 0.07 |  |  | 0.79 | 0.20 | 0.01 |
| comp45287_c1_seq1_139 | 0.91 | 0.09 |  |  | 0.74 | 0.25 | 0.01 |
| comp49053_c0_seq1_1206 | 0.94 | 0.06 |  |  | 0.74 | 0.25 | 0.01 |
| comp50406_c0_seq1_2537 | 0.30 | 0.50 | 0.20 |  | 0.19 | 0.70 | 0.11 |
| comp51904_c0_seq1_501 | 0.16 | 0.84 |  |  | 0.04 | 0.92 | 0.04 |

**Table S15a. Putative function of proteins containing SNPs significantly associated with budburst (BB2011). Function groups: RtE, response to environment; G&D, growth and development; R, reproduction. Where function is only linked to cellular processes, no function is indicated. Peptide ID and protein description obtained from uniprotkb_viridiplantae database. MAF: minor allele frequency (MAF < 0.05, rare; MAF > 0.05, common). Species: MU, *P. mugo*; SY, *P. sylvestris*; UN, *P. uncinata*. Analysis approach used: MLM, mixed linear model; MLMM, multi-locus mixed model.**

|  |  |  |  |  | Function group | | |  |
| --- | --- | --- | --- | --- | --- | --- | --- | --- |
| MAF | SNP | Species | Peptide ID | Protein | RtE | G&D | R | Reference |
| Analysis approach used: MLM | | | | | | | | |
| Common | comp51290_c0_seq5_216 | MU-UN MU-SY-UN | A9NP75 | AB hydrolase-1 domain-containing protein |  | x | x | Mindrebo et al 2016 |
|  | comp46858_c0_seq1_1024 | MU-UN MU-SY-UN | A0A2K1XXG8 | Uncharacterized protein [RNase T2 family] |  |  |  | Nurnberger et al 1990 |
|  | comp42130_c0_seq1_1318 | MU-SY-UN/ SY | A0A0D6R142 | DUF2470 domain-containing protein |  |  |  | Jung et al 2010 |
|  | comp47733_c0_seq1_4708 | MU-SY-UN/ SY | A0A2R6XJU9 | Sister chromatid cohesion protein |  |  |  | Peters and Nishiyama 2012 |
|  | comp46049_c0_seq1_1689 | MU-SY-UN/ SY | A0A0D6QZ13 | HTH myb-type domain-containing protein | x | x |  | Reviewed in Ambawat et al 2013 |
|  | comp52159_c0_seq1_436 | MU-UN | A0A3Q8BPZ8 | AUGMIN subunit3 |  |  |  | Goshima et al 2008 |
|  | comp48008_c0_seq1_836 | MU-UN | U5D3C7 | F-box domain-containing protein |  | x | x | Kipreos and Pagano 2000 |
|  | comp43117_c0_seq1_203 | MU-UN | D5A9S3 | Uncharacterized protein [Lipase 3 domain] | x |  |  | Shah 2005 |
|  | comp29799_c0_seq1_84 | MU-UN | A9NRE1 | Uncharacterized protein [Myb-like domain] |  |  |  | Klempnauer and Sippel 1987 |
|  | comp42558_c0_seq1_515 | MU-UN | A9NUZ9 | Magnesium-protoporphyrin IX monomethyl ester (oxidative) cyclase |  | x |  | Yang et al 2015; Kong et al 2016 |
|  | comp51726_c0_seq1_4360 | MU-UN | A0A200QX50 | Zinc finger protein | x | x | x | Takatsuji 1998 |
|  | comp18631_c0_seq1_563 | MU-UN | D5A976 | PMR5N domain-containing protein | x |  |  | Anantharaman 2010 |
|  | comp41446_c0_seq1_714 | SY | B8LQ84 | Long-chain-alcohol oxidase | x |  |  | Zhao et al 2008 |
|  | comp42661_c0_seq1_613 | SY | A0A200R3K9 | Rhodanese-like domain |  |  |  | Papenbrock et al 2010 |
|  | comp47733_c0_seq1_5033 | SY | A0A2R6XJU9 | Sister chromatid cohesion protein |  |  |  | Peters and Nishiyama 2012 |
|  | comp47733_c0_seq1_5101 | SY | A0A2R6XJU9 | Sister chromatid cohesion protein |  |  |  | Peters and Nishiyama 2012 |
|  | comp53584_c0_seq1_2201 | SY | B8LQR1 | Trehalase | x |  |  | Barraza and Sanchez 2013 |
|  | comp19484_c0_seq1_274 | SY | B8LLV2 | Uncharacterized protein [Helicase ATP binding domain] |  | x |  | Tuteja 2003 |
| Rare | comp48335_c0_seq3_629 | MU-UN | C0PQD2 | Dienelactone hydrolase domain-containing protein |  |  |  | Schlomann et al 1993 |
|  | comp50525_c0_seq1_2688 | MU-UN | A0A443N186 | DNA repair helicase XPD isoform X1 |  |  |  | Mindrebo et al 2016 |
|  | Doth_comp54682_c0_seq1_159_1118 | SY  MU-SY-UN | A0A5J5ADK7 | Fe2OG dioxygenase domain-containing protein |  | x |  | Sun et al 2020 |
|  | comp53528_c0_seq1_2969 | MU-UN | A0A3S3QE17 | Flowering time control protein FPA isoform X1 |  | x | x | Schomburg et al 2001 |
|  | comp53414_c0_seq1_1227 | MU-UN  MU-SY-UN | A9NZF1 | Gfo/Idh/MocA domain-containing protein |  |  |  | Taberman et al 2016 |
|  | comp54202_c0_seq2_791 | MU-UN | B8LS21 | Glycosyltransferase (EC 2.4.1.-) | x |  |  | Reviewed Keegstra and Raikhel 2001 and Gachon et al 2005 |
|  | comp48745_c0_seq3_659 | MU-UN MU-SY-UN | A0A453LKL4 | Histone H4 | x | x | x | Ding et al 2012; Song et al 2015; Zhu et al 2008 |
|  | comp51722_c0_seq1_418 | MU-UN | W1PTQ9 | HTH myb-type domain-containing protein | x |  |  | Ambawat et al 2013; Liu et al 2015; Wang et al 2015 |
|  | comp54389_c0_seq1_1101 | MU-UN MU-SY-UN | A0A199V6C0 | LMBR1 domain-containing protein A |  |  |  | No known function in plants |
|  | comp26730_c0_seq1_2554 | SY | A0A2H3Y3P8 | N-terminal acetyltransferase A complex auxiliary subunit NAA15-like |  | x | x | Feng et al 2016 |
|  | comp39509_c0_seq1_5557 | MU-UN | A0A1U8A2S8 | Paired amphipathic helix protein Sin3-like 4 isoform X1 | x | x | x | Huang et al 2019 |
|  | comp37834_c0_seq1_118 | MU-UN MU-SY-UN | A0A443NIB8 | Pentatricopeptide repeat-containing protein | x | x |  | Reviewed in Barkan and Small 2014 |
|  | comp31899_c0_seq1_3609 | MU-UN MU-SY-UN | A0A1U8A204 | Phospholipid-transporting ATPase (EC 7.6.2.1) | x |  |  | Chaffai and Koyama 2011; Underwood et al 2017; Poulsen et al 2008 |
|  | comp19208_c0_seq1_2090 | SY | A9NWK0 | Protein disulfide-isomerase (EC 5.3.4.1) |  |  |  | Houston et al 2005 |
|  | comp55695_c0_seq3_1742 | MU-UN | A0A199UVT1 | Putative U5 small nuclear ribonucleoprotein 200 kDa helicase |  | x | x | Hanley and Schuler 1991 |
|  | comp53226_c0_seq1_1405 | SY  MU-SY-UN | D5ABG3 | RING-type domain-containing protein | x | x |  | Wu et al 2014; Matsuda et al 2001; Von Armin and Deng 1993; Pepper and Chory 1997; Schumann et al 2003; Wang et al 2006; Zhang et al 2007 |
|  | comp49247_c0_seq1_1006 | MU-UN | A9NNU0 | Small heat shock proteins [sHSP] domain-containing protein | x |  |  | Waters 2012; Sun et al 2002 |
|  | comp52794_c0_seq1_2165 | SY | A0A2H3YGP1 | SNARE-interacting protein KEULE |  | x |  | Wu et al 2013 |
|  | comp49205_c0_seq1_2527 | MU-UN MU-SY-UN | B8LKN3 | Sulfate transporter and anti-sigma) STAS domain-containing protein |  |  |  | Sibagaki and Grossman 2004 |
|  | comp53736_c0_seq1_314 | MU-UN MU-SY-UN | A9P0D6 | TauD domain-containing protein |  |  |  | Unknown function in plants |
|  | comp42063_c0_seq1_922 | MU-UN MU-SY-UN | D5A7U4 | U1 small nuclear ribonucleoprotein C [U1C] |  | x |  | Golovkin and Reddy 2003 |
|  | comp55130_c0_seq108_549 | SY  MU-SY-UN | A0A0D6QWA0 | Uncharacterized protein [glycosyl hydrolase family] | x | x |  | Xu et al 2004; Minic 2008 |
|  | comp52277_c0_seq1_1396 | MU-UN MU-SY-UN | A0A2R6WET2 | Uncharacterized protein [Protein kinase domain] | x |  |  | Stone and Walker 1995 |
|  | comp52760_c0_seq1_196 | MU-UN MU-SY-UN | A9NWA3 | Uncharacterized protein [Protein kinase domain] | x |  |  | Stone and Walker 1995 |
|  | comp19471_c0_seq1_746 | MU-UN MU-SY-UN | W1PUS8 | Uncharacterized protein [putative function: Rab GTPase binding] | x | x |  | Kwon et al 2009; reviewed in Rutherford and Moore 2002 |
|  | comp48223_c0_seq1_312 | MU-UN | A0A443N803 | WD40 repeat | x | x |  | Stirnimann et al., 2010; Mishra et al 2012; Bashline et al 2015 |
|  | comp48223_c0_seq1_5041 | MU-UN MU-SY-UN | A0A443N803 | WD40 repeat | x | x |  | Stirnimann et al., 2010; Mishra et al 2012; Bashline et al 2015 |
|  | comp48223_c0_seq1_688 | MU-UN MU-SY-UN | A0A443N803 | WD40 repeat | x | x |  | Stirnimann et al., 2010; Mishra et al 2012; Bashline et al 2015 |
|  | comp45269_c0_seq1_760 | SY | A0A200RE06 | Xyloglucan endotransglucosylase/hydrolase (EC 2.4.1.207) | x | x | x | Cho et al 2006; Choi et al 2011; Van Sandt et al 2007; Hyodo et al 2003; Kurasawa et al 2009; Miedes et al 2013; |
|  | comp58475_c0_seq1_843 | MU-UN | A0A0D6R8S1 | J domain-containing protein |  |  |  | Miernyk et al 2001 |
| Analysis approach used: MLMM | | | | | | | | |
| Rare | comp31899_c0_seq1_3609 | MU-UN | A0A1U8A204 | Phospholipid-transporting ATPase | x |  |  | Chaffai and Koyama 2011; Underwood et al 2017; Poulsen et al 2008 |
|  | comp48223_c0_seq1_312 |  | A0A443N803 | Trp-Asp (WD40) repeats signature | x | x |  | van Nocker and Ludwig 2003; Mishra et al 2012; Lee et al 2010; Zhu et al 2008, Ramsay and Glover (2005), Zeng et al 2009 |
|  | comp53528_c0_seq1_2969 |  | A0A3S3QE17 | Flowering time control protein FPA isoform X1 | x |  | x | Schomburg et al 2001 |
|  | comp55037_c0_seq11_249 |  | B8LRT5 | Epimerase domain-containing protein | x | x |  | Rosti et al 2007 |
|  | comp45269_c0_seq1_760 | SY | A0A200RE06 | Xyloglucan endotransglucosylase/hydrolase |  | x |  | Miedes et al 2013 |
|  | comp49635_c0_seq1_343 |  | C0PQ40 | Uncharacterized protein [Membralin family] |  | x |  | Shimada et al 2020 |
|  | comp52794_c0_seq1_2165 |  | A0A2H3YGP1 | SNARE-interacting protein KEULE |  | x |  | Wu et al 2013 |
|  | comp31899_c0_seq1_3609 | SY-UN-MU | A0A1U8A204 | Phospholipid-transporting ATPase | x |  |  | Chaffai and Koyama 2011; Underwood et al 2017; Poulsen et al 2008 |
|  | comp53226_c0_seq1_1405 |  | D5ABG3 | RING-type domain-containing protein | x |  |  | Wu et al 2014; Matsuda et al 2001; Von Armin and Deng 1993; Pepper and Chory 1997; Schumann et al 2003; Wang et al 2006; Zhang et al 2007 |
|  | comp55037_c0_seq11_249 |  | B8LRT5 | Epimerase domain-containing protein | x |  |  | Rosti et al 2007 |

**Table S15b. Putative function of proteins containing SNPs significantly associated with traits: bud set (BS2010). All SNPs identified using the mixed linear model (MLM) approach in the *P. sylvestris* (SY) dataset. Function groups: RtE, response to environment; G&D, growth and development; R, reproduction. Peptide ID and protein description obtained from uniprotkb_viridiplantae database. MAF: minor allele frequency (MAF < 0.05, rare; MAF > 0.05, common).**

|  |  |  | Function group | | |  |
| --- | --- | --- | --- | --- | --- | --- |
| SNP | Peptide ID | Protein | RtE | GD | R | Reference |
| *MAF: Common* |  |  |  |  |  |  |
| comp47469_c0_seq1_2446 | W1NYN1 | C2H2-type domain-containing protein |  | x |  | Huang et al 2004 |
| comp52528_c0_seq1_927 | A9NL37 | 60S ribosomal protein L36 |  | x |  | Byrne 2009 |
| comp33699_c0_seq1_1791 | A0A438KK07 | Plant intracellular Ras-group-related LRR protein 4 |  | x |  | Forsthoefel et al 2005 |
| comp22360_c0_seq1_1093 | B8LRF3 | Cysteine dioxygenase | x |  |  | White et al (2017) |
| comp45158_c0_seq1_1173 | A9P2D5 | PRAI [phosphoribosyl anthranilate isomerase] domain-containing protein |  | x |  | He and Li 2001 |
| *MAF: Rare* |  |  |  |  |  |  |
| comp44449_c0_seq1_63 | A9NX75 | Carboxypeptidase (EC 3.4.16.-) |  | x |  | Kawakatsu et al 2009; Lehfeldt et al 2000 |
| comp50197_c0_seq1_652 | A0A1U8A9C6 | CLIP-associated protein [CLASP]-like |  | x |  | Ambrose et al 2007; Kirik et al 2007 |
| comp53726_c0_seq1_54 | A0A1U7Z0B6 | Filament-like plant protein 4 |  | x |  | uniprot database |
| comp19815_c0_seq1_115 | A9NN04 | Inhibitor I9 domain-containing protein |  |  |  | Figueiredo et al 2017 |
| comp55659_c0_seq1_1054 | A0A2K3NEL7 | LRR receptor-like kinase resistance protein (Fragment) | x | x |  | Liu et al 2017 and Dievart and Clark 2004 |
| comp48583_c0_seq1_703 | A0A0D6R2X0 | PPM-type phosphatase domain-containing protein | x |  |  | Rodriguez 1998; Schweighofer et al 2004 |
| comp44180_c0_seq1_322 | A0A1D1Y2C4 | Protein TOPLESS |  | x |  | Szemenyei *et al*. 2008 |
| comp42043_c0_seq1_534 | D5ADA7 | TH1 domain-containing protein |  |  |  | Syamaladevi and Sowdhamini 2012 |
| comp46712_c0_seq1_269 | A0A443NZW2 | Thyroid adenoma-associated protein | x | x |  | Dong et al 2018 |
| comp20404_c0_seq1_397 | B8LMB0 | Uncharacterized protein [Bax inhibitor 1 related family] |  | x |  | Watanbe and Lam 2006 |
| comp49104_c0_seq1_528 | A0A0D6R4E1 | Universal stree protein [USP] domain-containing protein | x |  |  | Lee et al 2019; Isokpehi et al 2011 |

**Table S15c. Putative function of proteins containing SNPs significantly associated with traits: growth (height: H2011, H2012, H2013; increment: I2012, I2013). Function groups: RtE, response to environment; G&D, growth and development; R, reproduction. Peptide ID and protein description obtained from uniprotkb_viridiplantae database. MAF: minor allele frequency (MAF < 0.05, rare; MAF > 0.05, common). Species: MU, *P. mugo*; SY, *P. sylvestris*; UN, *P. uncinata*. Analysis approach used: MLM, mixed linear model; MLMM, multi-locus mixed model.**

|  |  |  |  |  | Function group | | |  | |
| --- | --- | --- | --- | --- | --- | --- | --- | --- | --- |
| MAF | SNP | Species | Peptide ID | Protein | RtE | GD | R | | Reference |
| *Analysis approach used: MLM* | | | | | | | | | |
| Common | comp40846_c0_seq1_2337 | MU-UN MU-SY-UN | A0A0D6R0F5 | Receptor-like serine/threonine-protein kinase | x |  |  | | Diedhiou et al 2008; Sun et al 2013 |
|  | comp41840_c0_seq1_158 | SY  MU-SY-UN | A0A0D6QYF7 | Uncharacterized protein [Ankryin repeat region and RING domain] | x |  |  | | Cao et al 1997 |
|  | comp44927_c0_seq1_1219 | MU-SY-UN | W1P290 | PPIase cyclophilin-type domain-containing protein |  | x |  | | Wang and Heitman 2005 |
|  | comp47383_c0_seq1_174 | MU-SY-UN | W9SKC9 | Putative serine/threonine-protein kinase | x |  |  | | Diedhiou et al 2008; Sun et al 2013 |
|  | comp47901_c0_seq1_977 | MU-SY-UN | A0A2R6XVX4 | Ethylene receptor | x | x |  | | Abeles et al 1992 |
|  | comp49053_c0_seq1_1206 | MU-UN MU-SY-UN | A0A2H3Y947 | mitochondrial arginine transporter BAC2 | x |  |  | | Planchais et al 2014 |
|  | comp50406_c0_seq1_2537 | MU-UN | W1P8F1 | Uncharacterized protein [Histone acetyltransferase HAC-like protein] |  | x | x | | Deng et al 2007 and Boycheva et al 2014 |
|  | comp50686_c0_seq1_570 | SY | A0A0D6QXJ1 | Exo84C domain-containing protein |  | x |  | | Reviewed in Du et al 2018 |
|  | comp51669_c0_seq5_124 | MU-SY-UN | D5A9S5 | RNA recognition motif (RRM) domain-containing protein |  | x | x | | Lorkovic 2009 |
|  | comp51904_c0_seq1_501 | MU-UN MU-SY-UN | A0A0G7ZNW4 | Putative caffeoyl-CoA O-methyltransferase | x |  |  | | Yang et al 2017 |
|  | comp53296_c0_seq2_1323 | MU-SY-UN | B8LMA1 | Uncharacterized protein (Thiolase superfamily] | x | x | x | | Pye et al 2010 and Jin et al 2012 |
|  | comp53606_c0_seq1_1256 | SY | A0A1U7ZWK3 | Leucine-rich repeat receptor-like protein kinase (LRR-RLK) TDR isoform X1 | x | x |  | | Liu et al 2017 and Dievart and Clark 2004 |
|  | comp57469_c0_seq1_1389 | SY  MU-SY-UN | A9NXP5 | Eukaryotic translation initiation factor 3 subunit H (eIF3h) |  | x |  | | Kim et al 2004 |
|  | comp78883_c0_seq1_932 | MU-SY-UN | A0A0D6R4J7 | Exonuclease domain-containing protein |  | x |  | | Mason and Cox 2012, Tang and Sakamoto 2011 |
| Rare | comp57875_c0_seq1_2681 | MU-SY-UN | A0A0C9RLE4 | RING-in-between-RING [RBR]-type E3 ubiquitin transferase (EC 2.3.2.31) |  |  |  | | Unknown function in plants |
|  | comp43635_c0_seq1_122 | SY | A0A1V1FUY4 | U-box protein | x | x | x | | Azevedo et al 2001; Stone et al 1999; Gonzalez-Lamothe et al 2006; Kinoshita et al 2015 |
|  | comp50796_c0_seq1_1297 | SY | A0A0A7M6A6 | 26S proteasome regulatory subunit | x | x | x | | Reviewed by Vierstra 2009 |
|  | comp19094_c0_seq1_1018 | SY | A0A248Y4Q8 | CONSTANS-LIKE [COL]16 |  | x |  | | Ohmiya et al 2019 |
|  | comp19814_c0_seq1_714 | SY | A0A0D6QT61 | Cyclic nucleotide-binding domain-containing protein |  |  |  | | Bridges et al 2005 |
|  | comp45673_c0_seq1_2716 | SY | A0A0D6R702 | GRAS domain-containing protein |  | x |  | | Bolle 2004 |
|  | comp39306_c0_seq1_355 | MU-UN MU-SY-UN | H9VWZ8 | MoaC domain-containing protein (Fragment) |  | x |  | | Fido et al 1977 |
|  | comp51704_c0_seq1_1790 | SY | A0A371F2J6 | NAC domain-containing protein 73 | x |  |  | | Puranik et al 2012 |
|  | comp47519_c0_seq1_2584 | SY | A0A0D6R2P1 | Peptidase_M48 domain-containing protein |  |  |  | | Bhuiyan and van Wijk 2017 |
|  | comp46058_c0_seq1_1011 | SY | A0A0D6QVG3 | PPM-type phosphatase domain-containing protein | x |  |  | | Rodriguez 1998; Schweighofer et al 2004 |
|  | comp33384_c0_seq1_1437 | SY | A0A2P5EJA1 | Protein phosphatase | x |  |  | | Rodriguez 1998; Schweighofer et al 2004 |
|  | comp47138_c0_seq1_290 | SY | A0A5B6YSQ0 | Putative DNA excision repair protein ERCC-1 (Fragment) |  |  |  | | Hays 2002 |
|  | comp58475_c0_seq1_843 | MU-UN | A0A0D6R8S1 | J domain-containing protein |  |  |  | | Miernyk et al 2001 |
| *Analysis approach used: MLMM* | | | | | | | | | |
| Rare | comp39306_c0_seq1_355 | MU-UN | H9VWZ8 | MoaC domain-containing protein |  | x |  | | Fido et al 1977 |
|  | comp31134_c0_seq1_289 | SY | A0A2G5CSS6 | SPX domain-containing protein |  |  |  | | Secco et al 2012 |
|  | comp44962_c0_seq1_395 |  | RL15B | 60S ribosomal protein L15-2 |  | x |  | | Bobik et al 2018 |
|  | comp47519_c0_seq1_2584 |  | A0A0D6R2P1 | Peptidase_M48 domain-containing protein |  | x |  | | Lundquist et al 2012 |
| Rare | comp45357_c0_seq1_346 | SY-UN-MU | A0A2X0S4Q6 | RNase H domain-containing protein |  |  |  | | Majorek et al 2014 |
| Common | comp46813_c0_seq1_315 |  | A0A0D6QR56 | Peptidase A1 domain-containing protein | x | x |  | | Simoes and Faro 2004; Mazorra-Manzano et al 2010 |

**Table S16. Basic statistics for each set of SNPs used in each predictive model, based on allele frequencies in the *P. sylvestris* glasshouse trial dataset**

|  | MAF: No | | | | |  | MAF: Yes | | | | |
| --- | --- | --- | --- | --- | --- | --- | --- | --- | --- | --- | --- |
| SNP set | N | H_O_ | H_S_ | F_ST_ | F_IS_ |  | N | H_O_ | H_S_ | F_ST_ | F_IS_ |
| All polymorphic SNPs | 15,019 | 0.17 | 0.16 | 0.06 | -0.06 |  | 7,712 | 0.32 | 0.30 | 0.06 | -0.07 |
| *Predictive models for budburst* | | | | | | | | | | | |
| Budburst (MU-SY-SY; MU-UN; SY) | 25 | 0.13 | 0.12 | 0.06 | -0.07 |  | 17 | 0.17 | 0.16 | 0.06 | -0.07 |
| Budburst (MU-SY-UN; SY) | 15 | 0.15 | 0.14 | 0.07 | -0.07 |  | 11 | 0.19 | 0.18 | 0.07 | -0.07 |
| Budburst (SY) | 13 | 0.15 | 0.14 | 0.03 | -0.07 |  | 9 | 0.20 | 0.19 | 0.03 | -0.07 |
| *Predictive models for growth* | | | | | | | | | | | |
| Growth (MU-SY-UN; SY) | 14 | 0.11 | 0.10 | 0.03 | -0.06 |  | 11 | 0.14 | 0.13 | 0.03 | -0.06 |
| Growth (SY) | 7 | 0.12 | 0.11 | 0.04 | -0.09 |  | 4 | 0.21 | 0.21 | 0.03 | -0.06 |

N: number of SNPs; H_O_: observed heterozygosity; H_S_: mean gene diversities within populations; F_ST_: population differentiation; F_IS_: inbreeding coefficient. MAF:No = No MAF filter applied; MAF:Yes = a MAF filter was applied to exclude all SNPs with MAF < 0.05
